# Reduced ER-mitochondria connectivity promotes neuroblastoma multidrug resistance

**DOI:** 10.1101/2021.03.01.433120

**Authors:** Jorida Coku, David M. Booth, Jan Skoda, Madison C. Pedrotty, Jennifer Vogel, Kangning Liu, Annette Vu, Erica L. Carpenter, Jamie C. Ye, Michelle A. Chen, Peter Dunbar, Elizabeth Scadden, Eiko Nakamaru-Ogiso, Yimei Li, Kelly C. Goldsmith, C. Patrick Reynolds, Gyorgy Hajnoczky, Michael D. Hogarty

## Abstract

Most cancer deaths result from progression of therapy resistant disease, yet our understanding of this phenotype is limited. Cancer therapies generate stress signals that act upon mitochondria to initiate apoptotic programs. We isolated mitochondria from neuroblastoma cell lines obtained from children at diagnosis and after relapse following failed therapy, and profiled responses to tBid and Bim, death effectors activated by therapeutic stress. Mitochondria from post-relapse models had markedly attenuated cytochrome c release (surrogate for apoptotic commitment) in comparison with patient-matched diagnostic models. Mitochondrial DNA content, size, and shape did not differ consistently. However, we used electron microscopy to identify reduced endoplasmic reticulum-mitochondria contacts (ERMCs) as correlated with therapy resistance. ERMCs form microdomains for the transfer of Ca^2+^ to mitochondria. We confirmed reduced Ca^2+^ transfer in resistant cells, with restoration by re-opposing ERMCs via genetically-encoded linkers. However, reduced Ca^2+^ transfer was not present in all ERMC-reduced cancers with therapy resistance, supporting Ca^2+^-independent mechanisms. Genetically or biochemically reducing ERMCs in therapy sensitive tumors phenocopied resistance, validating these inter-organelle contacts as physiologic regulators of apoptosis. Our work confirms the importance of ERMCs in stress signaling and provides a previously unrecognized mechanism for cancer cell resistance that is not exclusive to other contributors.

## INTRODUCTION

Over 600,000 people die of cancer each year in the US, most with progression of disease that is resistant to available treatments. Our understanding of the mechanisms underlying such broad resistance to diverse drug classes and therapeutic modalities remains limited, but includes altered drug transport into or out of the cancer cell, such as from increased activity of ATP-binding cassette transporters (1), and mutations in genotoxic response genes like *TP53* (2). The former has not been confirmed to contribute to resistance in cancers in vivo and does not explain resistance to drugs that are not substrates for such transporters, and the latter does not explain therapy resistance in tumors with retained p53 activity and DNA repair. More recently, the focus has shifted to studying resistance arising in response to inhibitors of oncogenic kinases, with secondary mutations in the drug-target, activation of bypass signals, and cellular plasticity identified as causal, yet even here a large proportion of acquired resistance remains unexplained and seeking additional contributors is warranted (3–8). In short, our incomplete understanding of the diverse contributors to therapy resistance remains a principal barrier to improving cancer outcomes.

We used the highly lethal childhood tumor, neuroblastoma, as a model system to investigate therapy resistance. Children with high-risk neuroblastoma are treated with intensive chemoradiotherapy with stem cell rescue, surgery, retinoid therapy and immunotherapy (9). Many patients have metastatic disease at the time of diagnosis yet have chemosensitive disease that responds to treatment with significant tumor regression, including complete responses. Despite this, half of all patients relapse with lethal therapy resistant disease (10–12). To study emergent therapy resistance, we established tumor cell lines from the same patients at diagnosis prior to treatment, and again at the time of relapse during or after treatment, providing near-isogenic tumor models in which post-relapse tumors demonstrate therapy resistance acquired in situ during exposure to multimodal therapy (10). Since mitochondria serve as platforms for integrating cellular stress and survival signals in real-time, largely governed by Bcl2 family interactions, we hypothesized they harbor information related to therapeutic stress sensitivity. We used an unbiased assay in which cancer mitochondria are probed with tBid protein or BimBH3 peptides, death effectors activated downstream of most therapeutic stress (13–16), to define their sensitivity for activating mitochondrial outer membrane permeabilization (MOMP) as a surrogate for apoptotic commitment. We discovered that mitochondria from therapy resistant tumors have markedly attenuated MOMP responses in comparison with matched therapy sensitive counterparts, demonstrating that mitochondrial apoptotic signaling dysfunction arises during the course of clinical therapy and contributes to emergent multidrug resistance.

Many mitochondrial functions, including sensitivity to apoptosis, are regulated by signals derived from endoplasmic reticulum (ER) at contact sites with mitochondria [reviewed in (17, 18)]. Endoplasmic reticulum mitochondria contacts (ERMCs) are supported by ER-mitochondrial tethers, and upon cell fractionation, give rise to mitochondria-associated ER membranes (MAMs) that are enriched for specialized protein bridges mediating inter-organelle communication. For simplicity, we use MAM to refer to both ERMCs and MAMs in this paper. How cells regulate these contacts is incompletely understood, however, the importance of MAMs in tuning the cross-talk between these dynamic organelle networks is proposed to contribute to many pathophysiologic states, such as diabetes, neurodegeneration, and cancer (19–22). Here, we show that post-relapse tumor cell mitochondria have reduced MAM numbers and increased gap-distances, and these alterations are direct contributors to their stress resistance, consistent with the multidrug resistance phenotype seen clinically. We show this novel resistance phenotype contributes to resistance to chemotherapy, radiotherapy and molecularly-targeted agents. Importantly, it is not exclusive to other resistance mechanisms, but by acting downstream at a terminal signaling node imbues cancer cells with resistance to diverse therapy-induced stressors that engage mitochondria. This new framework for understanding therapy resistance may provide opportunities to enhance cancer care, including the measurement of relative resistance by characterizing MAM abundance and proximity, and enabling interventions to restore ER-mitochondrial communication in resistant cancers.

## RESULTS

### Mitochondria from drug resistant tumor cells have attenuated apoptotic signal transduction

Optimizing approaches developed by Letai (23), we isolated mitochondria from paired neuroblastoma cell lines derived from the same patients at diagnosis (DX) before therapy, and at relapse (REL) during or after completion of therapy (Figure 1A). Mitochondria-enriched heavy membrane fractions from tumor cells were incubated with recombinant truncated-Bid or the death-activating BH3 domain peptide of Bim (tBid and BimBH3, respectively) across a range of concentrations. Bid and Bim proteins are direct-activators of intrinsic apoptosis liberated by diverse cell stressors to either engage Bak or Bax to induce MOMP and cell death, or be sequestered by pro-survival Bcl2-family proteins to neutralize their death signals (15, 24). The sensitivity for release of cytochrome c in response to tBid or BimBH3 reflects a cell’s proximity to its apoptotic threshold (16, 25). By delivering terminal death effectors directly to mitochondria, the assay bypasses the contributions of drug transport, metabolism, target engagement and transcriptional response. Instead, mitochondrial responses reflect the state of the Bcl2 family and related apoptosis-regulating processes present in the tumor cell at the time of testing [reviewed in (26)]. A Bid BH3 domain with substitution of two highly-conserved residues served as a negative control and induced release of <10% of available cytochrome c in all experiments.

**Figure 1.**
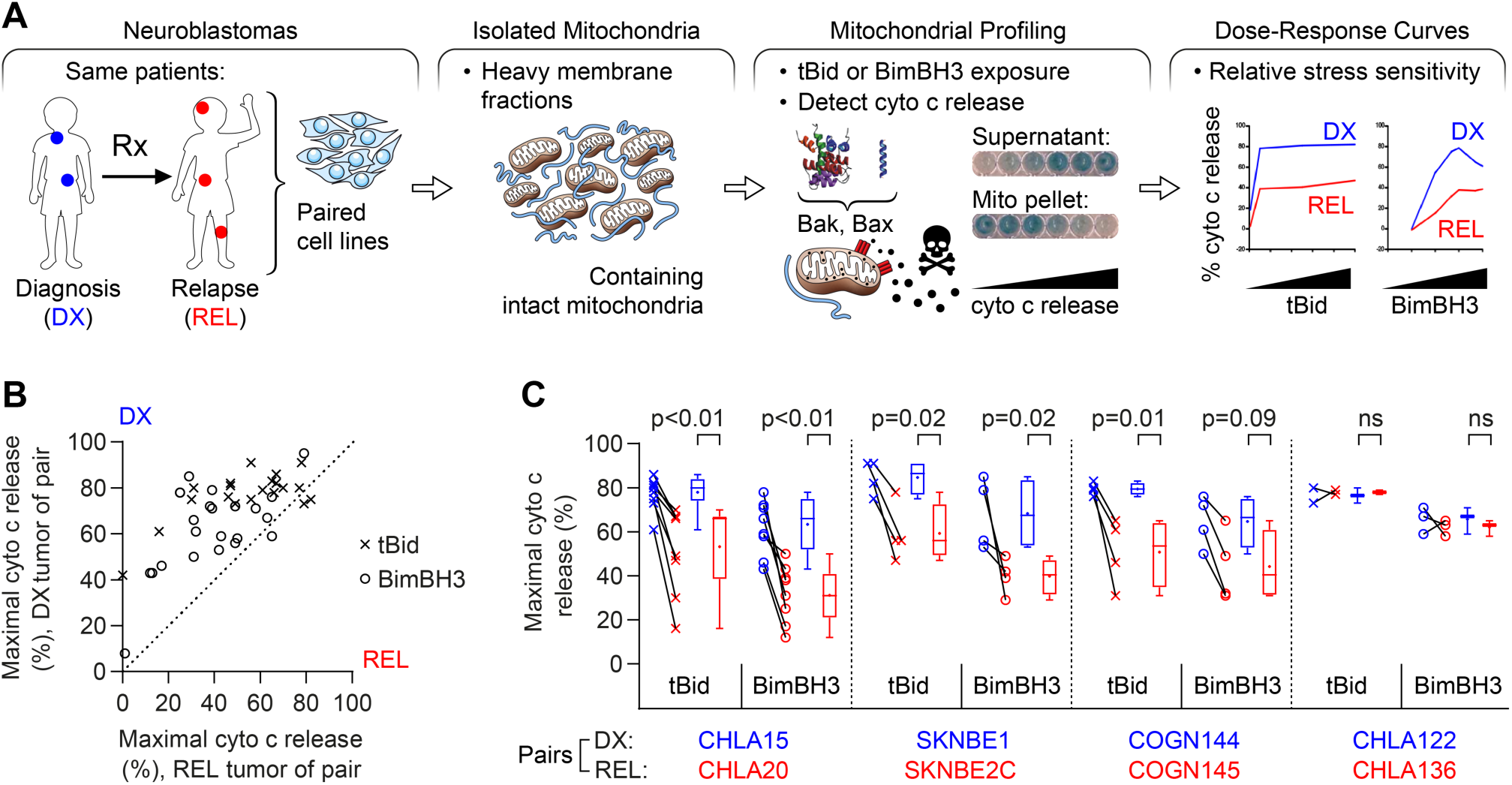
Mitochondria from REL neuroblastomas have attenuated apoptotic responses. **(A)** Tumor models were derived from the same children both at the time of diagnosis and at relapse following treatment. Mitochondria-enriched heavy membrane fractions were exposed to death-activating stimuli, tBid (1-500 nM) or BimBH3 peptide (1-10,000 nM). The proportion of cytochrome c released from mitochondria was measured as a surrogate for MOMP and apoptotic commitment. **(B)** Maximal cytochrome c release in response to tBid (x) or BimBH3 peptide (o), for each replicate of a DX/REL tumor pair, with at-diagnosis (DX) tumor cell release on the y-axis and post-relapse (REL) release on the x-axis. Each data-point is the mean of two technical replicates and all experiments are shown (1-9 biological replicates per DX/REL pair; n=44 total). **(C)** Maximal cytochrome c release for the DX/REL pairs with ≥3 biological replicates (DX= blue; REL= red; each experimental result connected by a line); box-whisker plots summarize data for each pair (box 25%-75%; belt=median; dot=mean; whiskers=minimum and maximum values). Statistical analyses were performed using an unpaired two-sided Student’s t-test, with significance p<0.05. Cyto c, cytochrome c; mito, mitochondria. Complete mitochondrial dose-response data set provided in Supplemental Figure 1.

Seven DX/REL matched cell line pairs were studied, each pair tested concurrently under identical culture conditions to minimize variability. Six of 7 pairs showed attenuated cytochrome c release from tumor cells derived at relapse (Figure 1B-C; Supplemental Figure 1A). For 5 of the 7, REL mitochondria had reduced cytochrome c release in response to both tBid and BimBH3 in every biological replicate, with both reduced sensitivity for release and reduced maximal release. One pair demonstrated reduced cytochrome c release in response to BimBH3 peptide but not tBid; and one pair (CHLA122/CHLA136) showed no consistent change as maximal release differed <10% in all but one replicate. Overall, in 41 of 44 (93%) assays with tBid or BimBH3, the mitochondria from post-relapse tumor cells released less cytochrome c than their patient-matched counterparts from the time of diagnosis (Figure 1B). Recombinant tBid was more potent at inducing cytochrome c release than BimBH3 peptide (all experiments), and maximal tBid-induced release exceeded that of BimBH3 in 39 of 42 (93%) experiments, despite BimBH3 peptide being used to >1 log higher concentrations.

Relative cytochrome c release from tumor cell mitochondria was reproducible (Figure 1C; Supplemental Figure 1). Mitochondria from CHLA15 (DX) and CHLA20 (REL) were profiled in 9 biological replicates with both tBid and BimBH3 and maximal cytochrome c release for CHLA15 was greater than CHLA20 in all 18 experiments. The SKNBE1(DX)/SKNBE2C(REL) and COGN144(DX)/COGN145(REL) pairs were profiled in 4 biological replicates each, and DX tumor mitochondria released more cytochrome c to both tBid and BimBH3 in all 16 experiments. In contrast, CHLA122 (DX) and CHLA136 (REL) showed nearly equivalent cytochrome c release in 4 of 5 experiments (in one, DX cells released >10% more cytochrome c than REL cells in response to BimBH3). In all, maximal cytochrome c release in response to tBid or BimBH3 peptide was significantly higher for DX tumor cells compared with patient-matched REL tumor cells for the CHLA15/CHLA20, SKNBE1/SKNBE2C and COGN144/COGN145 pairs, consistent with apoptosis resistance at relapse (Table 1). Overall, DX neuroblastomas released >50% of available cytochrome c in 38 of 43 of experiments (88%) using either tBid or BimBH3 as a stimulus, whereas REL neuroblastomas released >50% in just 19 of 44 of experiments (43%), further reflecting their attenuated response to stress (p<0.01). Only the CHLA122/CHLA136 pair demonstrated no difference in cytochrome c release in response to tBid or BimBH3.

**Table 1.** Summary of paired diagnostic (DX) and relapse (REL) neuroblastoma models.

|  | <u>CHLA15</u> | <u>CHLA20</u> |  | <u>SKNBE1</u> | <u>SKNBE2C</u> |  | <u>CHLA122</u> | <u>CHLA136</u> |  |
| --- | --- | --- | --- | --- | --- | --- | --- | --- | --- |
| **<u>MITOCHONDRIAL RESPONSE:</u>** |  |  |  |  |  |  |  |  |  |
| Max % cyto c release, tBid (mean +/- sd) | 78 ± 8 | 53 ± 19 | *p*<0.01 | 85 ± 7 | 59 ± 11 | *p*<0.02 | 77 ± 5 | 78 ± 1 | p= ns |
| Max % cyto c release, BimBH3 (mean +/- sd) | 63 ± 13 | 31 ± 12 | *p*<0.01 | 68 ± 14 | 40 ± 7 | *p*<0.02 | 66 ± 6 | 62 ± 4 | p= ns |
| **<u>TUMOR CELL RESPONSE (fold-resistance, IC<sub>50</sub>):</u>** |  |  |  |  |  |  |  |  |  |
| Mafofamide |  | 3.9 |  |  | 12.9 |  |  | 1.2 |  |
| Doxorubicin |  | 33.0 |  |  | 28.6 |  |  | 1.7 |  |
| Cisplatin |  | 1.6 |  |  | 4.2 |  |  | 2.6 |  |
| Etoposide |  | 3.5 |  |  | 122.0 |  |  | 2.1 |  |
| Ionizing Radiation |  | 2.1 |  |  | 2.0 |  |  | n.d. |  |
| Crizotinib |  | 5.0 |  |  | n.d. |  |  | n.d. |  |
| ABT737 |  | 44.0 |  |  | n.d. |  | 1.8 |  |  |
| **<u>MITOCHONDRIA:</u>** |  |  |  |  |  |  |  |  |  |
| Number analyzed | 196 | 241 |  | 137 | 160 |  | 206 | 143 |  |
| Perimeter, mean (nm) | 2,207 | 1,725 | *p*<0.01 | 1,653 | 1,810 | *p*=0.06 | 1,626 | 1,454 | p= ns |
| Perimeter, median (nm) | 2,055 | 1,675 |  | 1,503 | 1,725 |  | 1,404 | 1,330 |  |
| Circularity, median | 0.902 | 0.920 | *p*<0.01 | 0.910 | 0.898 | *p*= ns | 0.913 | 0.905 | p= ns |
| Roundness, median | 0.996 | 0.745 | *p*= ns | 0.714 | 0.692 | *p*= ns | 0.676 | 0.655 | p= ns |
| % with 0 MAMs | 18% | 32% | *p*<0.01 | 18% | 40% | *p*<0.01 | 17% | 8% | *p*<0.01 (REL fewer) |
| % with 1 MAMs | 39% | 39% |  | 37% | 35% |  | 33% | 34% |  |
| % with 2-3 MAMs | 41% | 25% | *p*<0.01 | 38% | 22% | *p*<0.01 | 42% | 43% | p= ns |
| % with ≥4 MAMs | 3% | 3% |  | 6% | 4% |  | 7% | 15% |  |
| %OMM with MAM gap-width ≤10 nm | 0.7% | 0.4% | (↓43%) | 2.3% | 1.1% | (↓52%) | 2.1% | 2.7% | (↑29%) |
| %OMM with MAM gap-width ≤25 nm | 2.5% | 2.1% | (↓16%) | 6.0% | 3.5% | (↓42%) | 6.1% | 6.8% | (↑11%) |
| %OMM with MAM gap-width ≤50 nm | 6.8% | 6.3% | (↓7%) | 12.4% | 7.6% | (↓39%) | 12.8% | 14.2% | (↑13%) |
| %OMM with MAM gap-width ≤100 nm | 16.9% | 15.5% | (↓8%) | 24.4% | 15.2% | (↓38%) | 24.0% | 25.5% | (↑6%) |
| **<u>MAM:</u>** |  |  |  |  |  |  |  |  |  |
| Number analyzed | 267 | 266 |  | 203 | 154 |  | 334 | 386 |  |
| MAMs per mitochondria | 1.36 | 1.10 | *p*<0.01 | 1.48 | 0.96 | *p*<0.01 | 1.62 | 2.00 | *p*=0.01 (REL more) |
| MAM frequency (per nm perimeter) | 1,651 | 1,556 |  | 1,116 | 1,880 |  | 710 | 1,134 |  |
| %MAM with gap-width ≤10 nm | 18% | 12% | *p*=0.05 | 30% | 21% | *p*<0.01 | 36% | 39% | p= ns |
| %MAM with gap-width ≤25 nm | 40% | 36% | *p*= ns | 57% | 46% | *p*<0.01 | 65% | 69% | p= ns |
| %MAM with gap-width ≤50 nm | 70% | 63% |  | 80% | 73% |  | 86% | 93% |  |
| %MAM with gap-width ≤100 nm | 100% | 100% |  | 100% | 100% |  | 100% | 100% |  |

### Cytochrome c release from isolated mitochondria correlates with tumor cell sensitivity to diverse therapeutic stressors

Attenuated mitochondrial responses to tBid and Bim, the stress sentinels activated by chemotherapy (13–16), predicts for reduced cytotoxicity in response to such drugs. We compared sensitivity to chemotherapeutics of different drug classes used to treat neuroblastoma, including cisplatin, mafosfamide, etoposide and doxorubicin. Etoposide and doxorubicin are substrates for the multidrug resistance protein, P-glycoprotein-1, whereas cisplatin and mafosfamide are not. Both the CHLA15/CHLA20 and SKNBE1/SKNBE2C pairs that had attenuated release of cytochrome c from REL mitochondria, showed relative chemoresistance in REL tumor cells to both P-glycoprotein-1 substrates and non-substrates (Figure 2A; Supplemental Figure 2A; Table 1). In contrast, the CHLA122/CHLA136 pair without attenuated mitochondrial release of cytochrome c did not show differential chemosensitivity. Fold-resistance by comparing IC_50_ values for DX/REL pairs were 1.6-fold to 33-fold for CHLA15/CHLA20 (3 of 4 drugs with >3-fold difference); 4.2-fold to 122-fold for SKNBE1/SKNBE2C; and 1.2-fold to 2.6-fold for CHLA122/CHLA136 (no drugs with >3-fold difference). We next assessed sensitivity to ionizing radiation as a treatment modality to which there is less cross-resistance than chemotherapy. Radiation bypasses drug transport and metabolism contributions to deliver genotoxic stress, which can be quantified as γ-H2AX foci. SKNBE2C (REL) cells had a larger induction of γ-H2AX foci post-radiation (∼10-fold compared with ∼5-fold for SKNBE1), yet they were 2-fold more radiation resistant (Supplemental Figure 3). However, SKNBE2C cells harbor an acquired *TP53* mutation (C135F) contributing to their resistance, and fail to induce p53, p21 or noxa. CHLA20 (REL) cells, however, are *TP53* wild-type and were also >2-fold radiation resistant in comparison with CHLA15 (DX) cells despite both inducing p53 response genes and deriving equivalent DNA damage, confirming attenuated apoptotic signaling downstream of radiation-induced genotoxic damage (Figure 2B-C).

**Figure 2.**
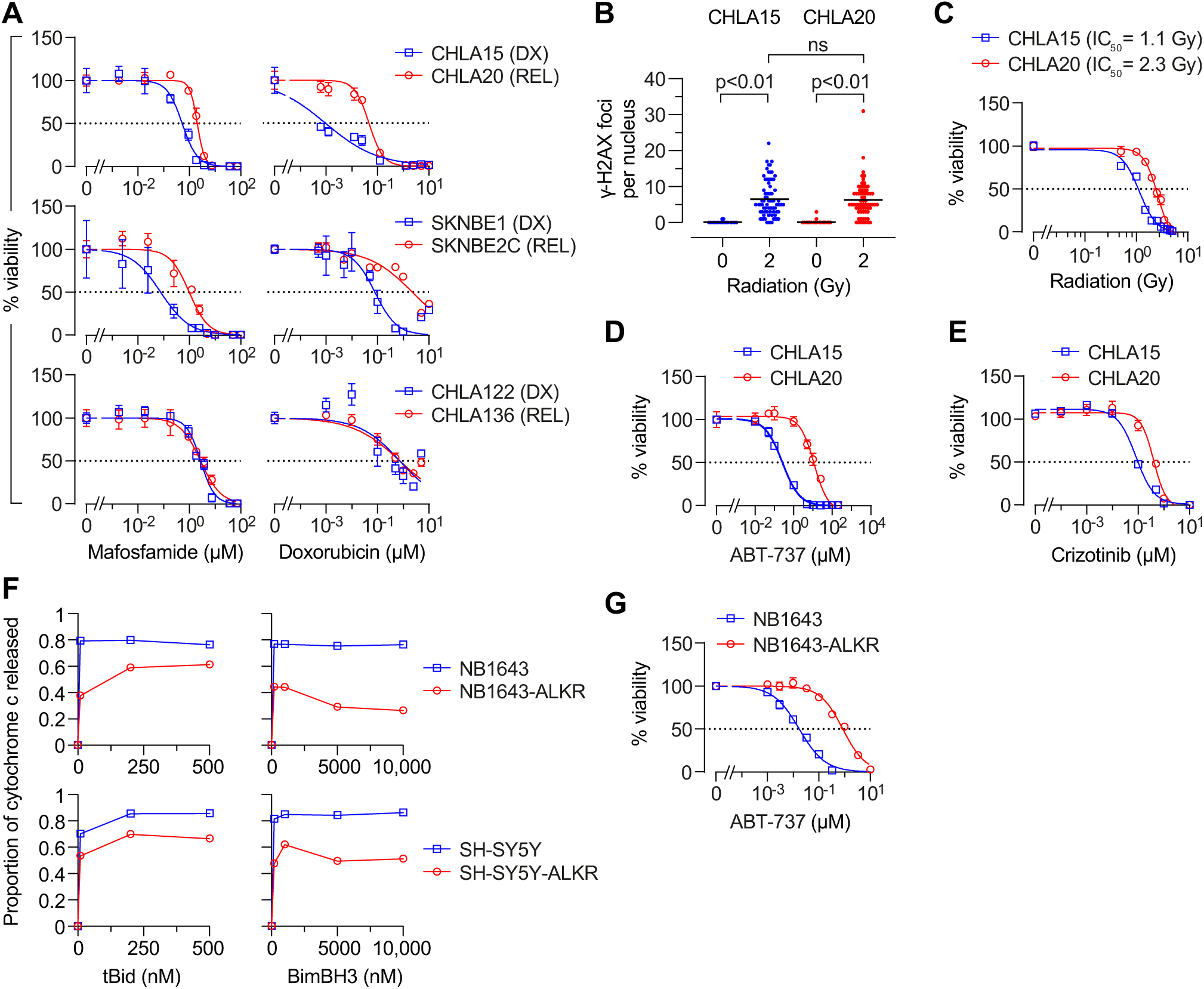
REL neuroblastomas are resistant to diverse cancer therapeutics. **(A)** In vitro viability curves measuring cellular ATP (Cell Titer-Glo) after 72hr exposure to mafosfamide or doxorubicin. Results are shown for three DX/REL neuroblastoma cell line pairs, two with attenuated MOMP in cytochrome c release studies (CHLA15/CHLA20 and SKNBE1/SKNBE2C) and one without (CHLA122/CHLA136). Data for cisplatin and etoposide are in Supplemental Figure S2A. **(B)** DNA-damage induced by 2 Gy ionizing radiation as measured by γ-H2AX foci at 1hr (n=53-77 cell nuclei/condition scored; mean shown); statistical analyses were performed using an unpaired two-sided Student’s t-test. In vitro viability for CHLA15 (DX) and CHLA20 (REL) following ionizing radiation at 7 days **(C)**, following 48hr exposure to the Bcl2/Bclx-inhibitor, ABT-737 **(D)**, or 120hr exposure to the Alk inhibitor, crizotinib **(E)**. **(F)** Cytochrome c release from mitochondria after exposure to tBid or BimBH3 peptide for parental NB1643 and SY5Y cells, in comparison to cells cultured in escalating concentrations of crizotinib until resistant (NB1643-ALKR and SY5Y-ALKR); complete data are provided in supplemental Figure S1B. **(G)** In vitro viability of NB1643 and NB1643-ALKR cells following 72hr exposure to ABT-737. DX, diagnosis; REL, relapse. For A, C-E, and G: data points are mean and SD from triplicate wells, experiments are representative of at least three biological replicates. For F, data points are mean of duplicate wells (SD<0.05 at all points) in a representative experiment from at least two biological replicates.

To determine whether attenuated mitochondrial responses also contribute to resistance to molecularly targeted drugs, we tested Bcl2 inhibitors since their mechanism of activity is localized at mitochondria. We previously showed that CHLA15 (DX) cells are sensitive to Bcl2 inhibitors as they use Bcl2 to sequester Bim and prevent its activation of Bak or Bax (25). Both CHLA15 (DX) and CHLA20 (REL) cells have Bim sequestered by Bcl2 and the Bcl2/Bclx inhibitor ABT-737 displaces Bim with equal potency, yet CHLA20 cells are >40-fold more resistant despite the patient the cell line derived from never having been treated with a Bcl2 inhibitor (IC_50_: CHLA15=260nM, CHLA20=11.4μM; Figure 2D). In contrast, the CHLA122/CHLA136 pair also have Bim sequestered by Bcl2, but these cells have similar mitochondrial responses and Bcl2 inhibitor responses (∼1.8-fold IC_50_ difference). We did not test SKNBE1/SKNBE2C cells with ABT-737 as they use Mcl1, rather than Bcl2, to neutralize Bim (25). We next studied anaplastic lymphoma kinase (Alk) inhibitors since mutations in the *ALK* gene are found in 10-14% of neuroblastomas and Alk inhibition has anti-tumor activity (27). Both CHLA15 (DX) and CHLA20 (REL) cells harbor an *ALK* R1275Q mutation (variant allele frequency of 0.46 and 0.49, respectively). The patient this tumor pair was obtained from had not been treated with an Alk inhibitor, yet REL cells were >3-fold more resistant to the Alk inhibitors crizotinib, ceritinib and lorlatinib, without having a secondary resistance-encoding *ALK* mutation (e.g., crizotinib IC_50_: CHLA15=96nM, CHLA20=349nM; Figure 2E). SKNBE1/SKNBE2C and CHLA122/CHLA136 have wild-type *ALK* and are resistant to Alk inhibitors (Supplemental Figure 2B). Therefore, attenuated mitochondrial signaling that arises in response to multimodal therapy in situ confers resistance to chemotherapy, radiotherapy, and molecularly-targeted drugs (the latter in the absence of prior selective pressure). We next assessed the reverse: whether selection for resistance to a molecularly targeted agent can induce this mitochondrial phenotype. We exposed SH-SY5Y (*ALK* F1174L) and NB1643 (*ALK* R1275Q) cells to escalating concentrations of crizotinib to generate crizotinib-resistant clones, and both demonstrated attenuated MOMP responses to tBid and BimBH3 in the absence of secondary *ALK* mutations, phenocopying REL cells (Figure 2F). In addition, such Alk inhibitor resistant cells were more resistant to chemotherapy drugs such as etoposide (>20-fold) and molecularly-targeted drugs like Bcl2 inhibitors (>40-fold; Figure 2G).

### Attenuated mitochondrial apoptotic responses are accompanied by reductions in ER-mitochondria associated membranes

We previously showed that loss of Bak or Bax was unlikely to be a driver of attenuated MOMP as DX and REL cells express abundant Bak and Bax protein (25). We next compared mitochondrial mass, size, shape and mitochondrial DNA (mtDNA) content from DX and REL pairs, as these have been correlated with apoptotic signaling in other systems. Mitochondrial mass assessed by citrate synthase activity was unchanged in CHLA15 and CHLA20, and reduced in SKNBE2C (relative to SKNBE1) and CHLA136 (relative to CHLA122; Supplemental Figure 4A). Mitochondrial DNA content was measured using quantitative-PCR for two mitochondrial genes (*MT-CO1* and *MT-ATP6*), each normalized to two nuclear genes (*CFAP410* and *MTTP*, disomic in >85% of neuroblastomas). *TP53* mutant SKNBE2C cells had markedly reduced mtDNA content compared with *TP53* wild-type SKNBE1 cells, consistent with p53 mutation effects on mtDNA content (28). CHLA15/CHLA20 and CHLA122/CHLA136 pairs had less divergent mtDNA quantity with modest reductions in REL cells at a subset of loci (Supplemental Figure 4B). We next analyzed transmission electron micrographs to assess mitochondrial size (circumference) and shape (roundness and circularity; Figure 3A-C). These did not differ for the SKNBE1/SKNBE2C and CHLA122/CHLA136 pairs, though there was a trend toward larger mitochondria in SKNBE2C cells (p=0.06). The CHLA15 (DX) and CHLA20 (REL) pair did differ, with REL cells having smaller more circular mitochondria (p<0.01). Overall, these features did not correlate with mitochondrial cytochrome c release sensitivity.

**Figure 3.**
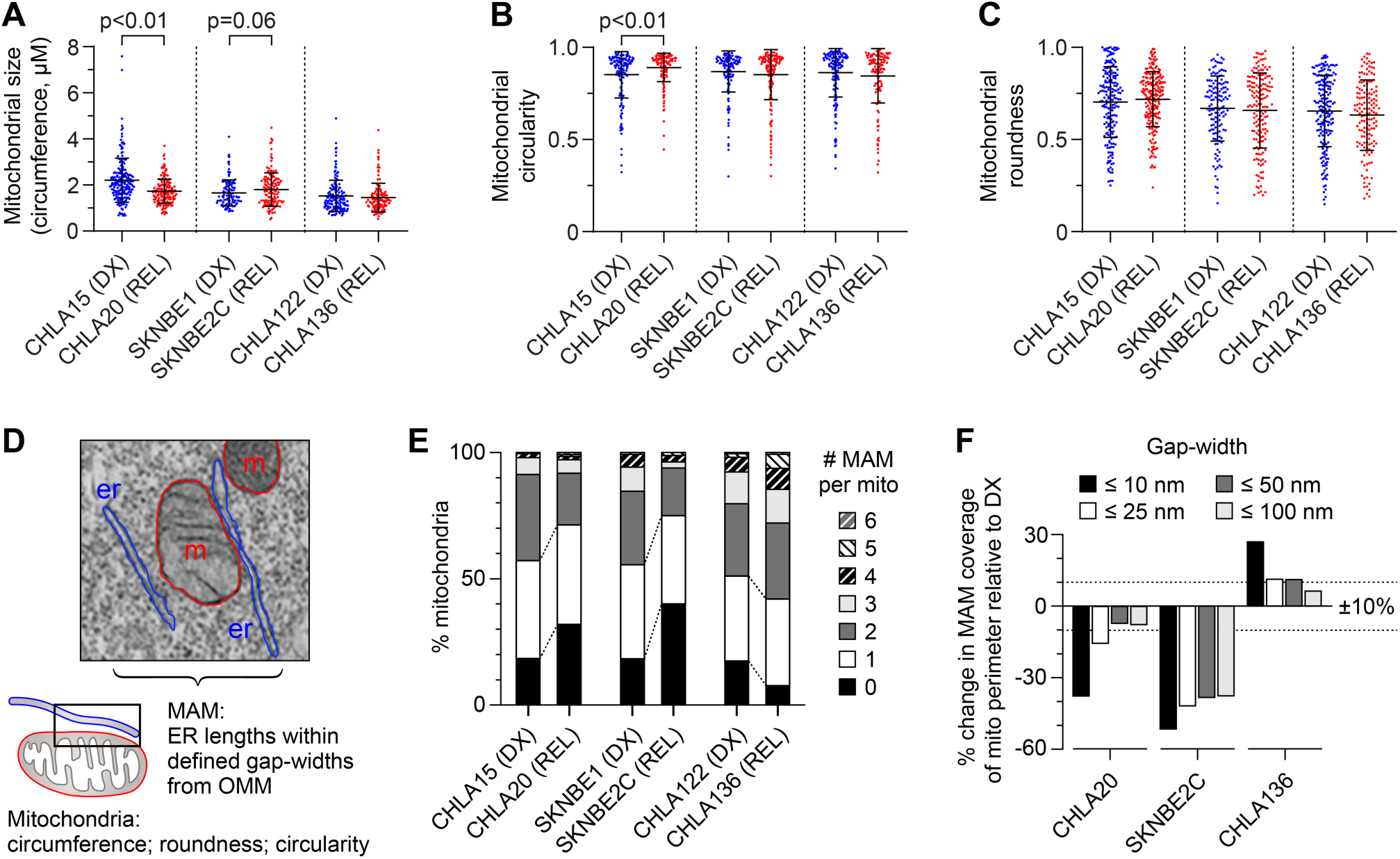
REL neuroblastomas have reduced mitochondria-associated membranes compared with patient-matched tumors from DX. Transmission electron microscopy (TEM) image analysis was used to quantify mitochondrial size **(A)**, circularity **(B)** and roundness **(C)** in DX/REL neuroblastoma pairs (mean +/-SD shown). **(D)** Electron micrograph illustrating organelle masking and characterization of MAM interface metrics extracted using an Image J custom macro (http://sites.imagej.net/MitoCare/). **(E)** Proportion of mitochondria with 0 to 6 or more MAMs are shown for each cell line, as DX/REL pairs. **(F)** The percentage of the mitochondria perimeter with an apposed ER within defined gap-widths (as shown) for DX/REL neuroblastoma cell line pairs. For A-C and E-F, n=137-241 mitochondrial per cell line. DX, diagnosis; REL, relapse; er, endoplasmic reticulum; m, mitochondria. Statistical analyses were performed using an unpaired two-sided Student’s t-test, with significance p<0.05 (trend, p<0.10).

We used electron microscopy to visualize the mitochondria-enriched heavy-membrane fractions tested in our mitochondrial profiling assays. Non-mitochondria organellar membranes were abundant in all yet there was an increase in the proportion of mitochondria in fractions from REL cells, despite their reduced cytochrome c release when stimulated (Supplemental Figure 5A-B). ER is a major contributor to heavy-membrane fractions, in particular, MAMs. We postulated that fewer ER-mitochondria contacts leads to reduced MAM content in heavy-membrane fractions. To test this, we fractionated heavy-membranes to resolve purified mitochondria from MAMs (29). With equal cellular input, more MAMs were visualized from DX cells than REL cells for the CHLA15/CHLA20 and SKNBE1/SKNBE2C pairs, while the CHLA122/CHLA136 pair that lacks attenuated mitochondrial signaling had slightly more MAM content in REL cells (Supplemental Figure 5C). Consistent with this, Chipuk et al have shown in murine liver and HeLa cells that separating mitochondria from MAMs using limiting proteolysis reduces their release of cytochrome c in response to Bid and BimBH3 domains (30). We also assessed ER-specific proteins in heavy-membrane fractions from DX and REL cells but were unable to detect differential expression (Supplemental Figure 5D).

We next directly visualized MAM interfaces using transmission electron micrographs of DX and REL tumor cells. We masked mitochondria and ER perimeters and defined MAMs as regions of ER within 100 nm of the outer mitochondrial membrane (OMM), and characterized their number, length and gap-width from the OMM, binned as ≤10 nm, 10-25 nm, 25-50 nm, and 50-100 nm (Figure 3D). CHLA20 (REL) cells had reduced MAM content compared with CHLA15 (DX) cells (Table 1). The number of MAMs per mitochondria was reduced (p<0.01), the frequency of mitochondria with ≥2 MAMs was reduced while those absent MAMs were increased (both p<0.01; Figure 3E; Table 1). Because CHLA20 mitochondrial had a mean circumference 22% smaller (p<0.01), MAMs occurred at equal frequency in DX/REL cells: every 1,651nm in CHLA15 and every 1,556nm in CHLA20. However, there was a marked reduction in closely-apposed MAMs and MAM lengths in REL cells. The proportion of MAM that came within 10nm of the OMM was reduced (p=0.05), as was the proportion of the OMM perimeter in contact with a MAM at <10nm (Figure 3F; Table 1). In contrast, the proportion approximated by a MAM at larger gap-distances were less reduced (Figure 3F). Similarly, SKNBE2C (REL) cells were reduced in the number of MAMs per mitochondria compared to SKNBE1 (DX) cells (p<0.01), and the frequency of mitochondria with ≥2 MAMs was reduced, while those absent MAMs were increased >2-fold (both p<0.01; Figure 3E; Table 1). MAMs occurred on average every 1,116 nm along the OMM of SKNBE1 but only every 1,880 nm in SKNBE2C. In addition to reduced frequency, the total length and proximity of MAMs were reduced in SKNBE2C cells, adjacent to 24% of the mitochondrial perimeter in SKNBE1 but only 15% in SKNBE2C cells, and MAMs were significantly reduced across all gap-widths (Figure 3F; Table 1).

In contrast, CHLA136 (REL) cells do not have attenuated mitochondrial responses, multidrug resistance, reduced MAM content by fractionation, nor reduced MAM interfaces in comparison with CHLA122 (DX) cells (Figure 3E-F; Table 1). In fact, CHLA136 (REL) cells had a trend toward more mitochondria with ≥2 MAMs (p=0.10) and fewer orphan mitochondria (p<0.01), more MAMs per mitochondria overall (p=0.01), and a slightly higher proportion of MAMs along the OMM across all gap-widths. The characteristics of individual MAMs by their length and relative proximity to the mitochondrial outer membrane were otherwise similar (Figure 3F). In all, reduced ER-mitochondria connectivity (in particular, a reduction in the proportion of MAMs in close proximity to the OMM) was recurrently present in tumor cells with multidrug resistance and attenuated mitochondrial responses to death stimuli, nominating this feature as a contributor to therapy resistance.

### Reduced transfer of Ca^2+^ from MAMs to mitochondria is not required for attenuated apoptotic signaling

MAMs support mitochondria as platforms to integrate stress signals. They facilitate transfer of calcium to mitochondria, which has been linked to Bcl2-family functions and apoptotic sensitivity (31, 32). ER provides the major intracellular reservoir for Ca^2+^, and MAMs are enriched for calcium release constituents to create localized microdomains to enable calcium transfer into mitochondria through the mitochondrial calcium uniporter (mtCU) that has low Ca^2+^ affinity (32). The ensuing rise in mitochondrial matrix calcium ([Ca^2+^]_m_) can promote apoptosis by either stimulating permeability transition pore opening in the inner membrane or facilitating Bak and/or Bax oligomerization and MOMP induction (33). Indeed, synthetically decreasing ER-mitochondria tethering in rat liver cells and basophils leads to reduced Ca^2+^ transfer and reduced apoptotic sensitivity, while augmenting tethering has been shown to increase both (34).

We posited that the reduction in MAMs in REL tumor cells reduces ER-to-mitochondria Ca^2+^ transfer to attenuate mitochondrial apoptotic signaling. To test this, we measured Ca^2+^ transfer in two DX/REL pairs with differential MAM content. Mitochondrial calcium concentration ([Ca^2+^]_m_) was detected with a fluorescent protein-based Ca^2+^ sensor targeted to the mitochondrial matrix, GCamp6f. Cytoplasmic calcium concentration ([Ca^2+^]_c_) was monitored with fura2 loaded to the cells as fura2AM. Calcium responses evoked by the IP_3_R-linked stimulus, carbachol, were recorded and the time-courses for the corresponding fluorescence signals calculated. CHLA20 (REL) cells have attenuated MOMP responses, multidrug resistant phenotype and reduced MAM content in comparison with CHLA15 cells (DX), yet both have similar [Ca^2+^]_c_ and [Ca^2+^]_m_ signals and coupling time (<2 seconds), the time difference between [Ca^2+^]_c_ and [Ca^2+^]_m_ achieving 50% of maximum (Figure 4A). Like CHLA20, SKNBE2C (REL) cells also have attenuated MOMP responses, multidrug resistant phenotype and reduced MAM content in comparison with SNKBE1 (DX) cells. However, unlike CHLA20, SKNBE2C cells have markedly reduced Ca^2+^ transfer compared with SKNBE1 (DX) cells, with a >3-fold increase in coupling time (p<0.01; Figure 4B). To confirm diminished Ca^2+^ transfer is due to a reduction in MAMs, we genetically augmented MAMs in SKNBE2C cells using a monomeric linker composed of a fluorescent protein extended with ER and OMM membrane anchor domains (34). Linker-induced augmentation of MAMs normalized the Ca^2+^ transfer coupling time to ∼1.5 seconds, confirming the reduction was a consequence of reduced ER-mitochondria contacts (Figure 4B). In both the CHLA15/CHLA20 and SKNBE1/SKNBE2C pairs the REL cells showed similar proportional reductions in MAM coverage at ≤10 nm ER-mitochondrial gap-width (Table 1). However, CHLA20 cells had preserved MAM coverage at larger gap-widths capable of accommodating the Ca^2+^ transfer machinery, while SKNBE2C cells were reduced at all gap-widths, limiting Ca^2+^ transfer.

**Figure 4.**
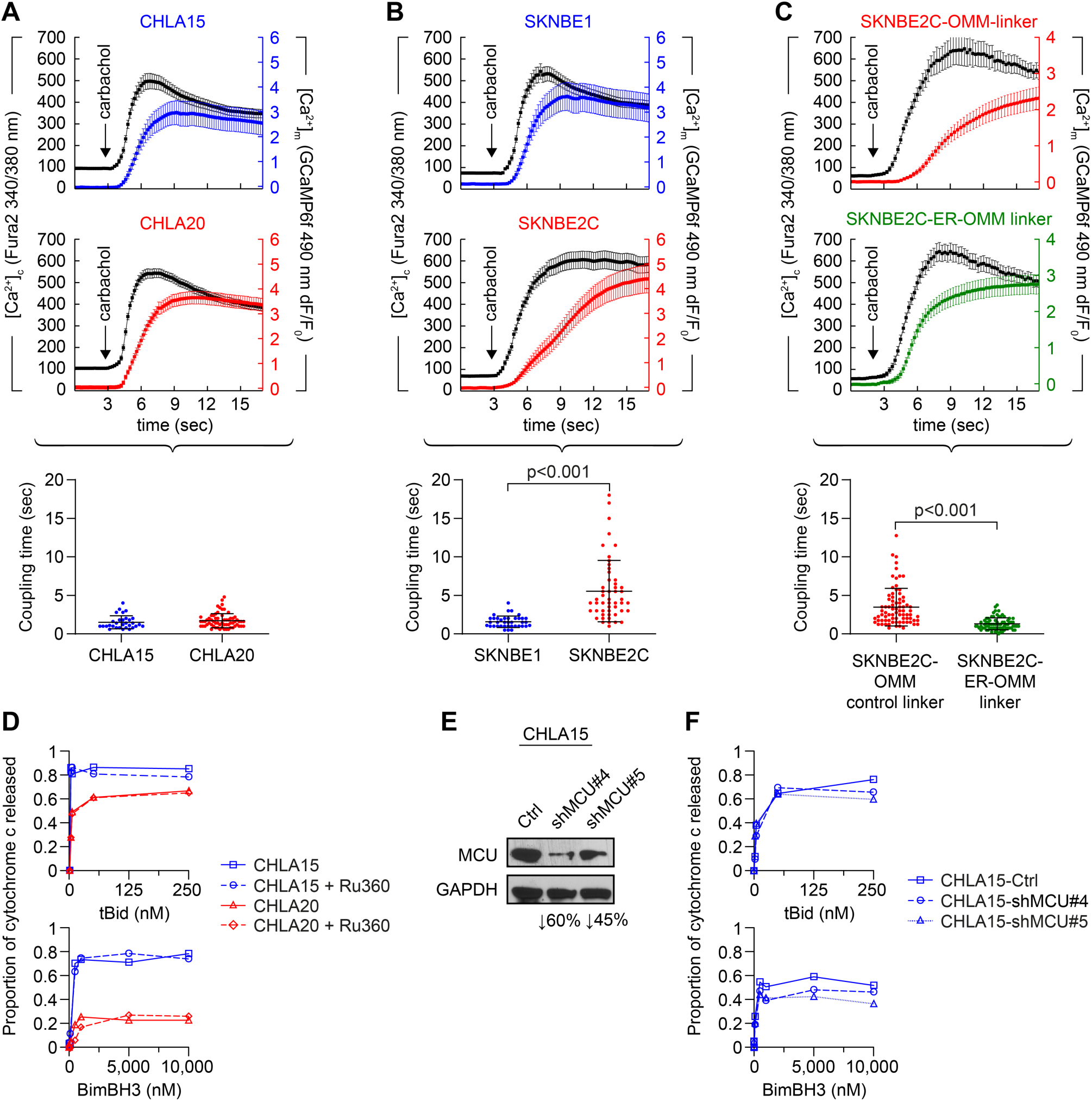
Reduced calcium transfer at ERMCs is not required for attenuated mitochondrial responses to therapeutic stress. Ca^2+^ transfer was assessed by measuring cytosolic (black-tracing) and mitochondrial (colored-tracing) Ca^2+^ concentration using fluorescent reporters as described in Methods. ER Ca^2+^ release was induced by 100μM carbachol, an inositol trisphosphate receptor (IP3R)-agonist; ∼time added indicated by arrow. Coupling time (time between achieving 50% of maximal cytosolic and mitochondrial calcium concentrations) is an index of ER-mitochondrial proximity and transfer efficiency. **(A-B)** Coupling time for two DX (blue)/REL (red) pairs with attenuated MOMP. **(C)** MAM proximity was enforced using an ER-OMM linker construct, with Ca^2+^ transfer coupling time shown for SKNBE2C cells expressing an OMM control linker, compared with those expressing an ER-OMM linker that enhances MAM proximity; mean +/- SD shown as whiskers. **(D)** Mitochondrial cytochrome c release for CHLA15 and CHLA20 cells following exposure to tBid or BimBH3 peptide, with or without Ru360 treatment. **(E)** Silencing of the mitochondrial calcium uniporter (MCU) was achieved in two CHLA15-shMCU clones, and cytochrome c release in response to tBid and BimBH3 assessed, **(F).** See also Supplemental Figure 6. OMM, outer mitochondrial membrane. For D and F, data points are mean of duplicate wells (SD<0.05 at all points) in a representative experiment from at least two biological replicates. Statistical analyses were performed using an unpaired two-sided Student’s t-test, with significance p<0.05 (trend, p<0.10). OMM, outer mitochondrial membrane.

We used biochemical and genetic approaches to further evaluate Ca^2+^ transfer and stress sensitivity. We treated CHLA15 heavy-membrane fractions with Ru360, a ruthenium red analog that inhibits the mtCU, before measuring cytochrome c release in response to tBid or BimBH3. No reduction in release was seen (Figure 4D), nor was cytochrome c release attenuated when treating whole cells short-term for 24 hours, or longer-term for 7 days, before obtaining mitochondrial profiles (data not shown). Similarly, CHLA20 (REL) cells did not have further attenuated cytochrome c release following exposure to Ru360, although their release was reduced in comparison with CHLA15, as expected. We next used siRNA to reduce expression of MCU, the pore-forming component of the mtCU (35, 36). We first showed that MCU was not differentially expressed in DX/REL pairs (Supplemental Figure 6A). MCU knockdown of ∼45% and 60% was achieved in two CHLA15 subclones and MOMP sensitivity was only modestly reduced (Figure 4E-F). We also tested MCU knock-down and Ru360 exposure in SKNBE1 cells and were unable to demonstrate a consistent reduction in cytochrome c release in response to tBid or BimBH3 (Supplemental Figure 6B-D). Collectively, these data support that reduced transfer of Ca^2+^ from ER to mitochondria is not required for mitochondrial desensitization towards apoptotic MOMP.

### The role of MAMs in mitochondrial apoptotic signaling can be functionally demonstrated

Limited proteolysis to cleave ER-mitochondria protein bridges led to reduced MOMP sensitization, however, it also induced >30% cytochrome c release under control conditions, dampening the dynamic range of the assay. Immunomagnetic-bead separation (30) induced less membrane disruption, and MOMP sensitivity in response to BimBH3 was greater for CHLA15 heavy membrane fractions (mitochondria with intact MAMs) compared with purified mitochondria (reduced MAMs; Figure 5A). SKNBE1 could not be assessed due to high release of cytochrome c release with either separation process. While it may be difficult to abolish MAMs by genetic knock-down of a single constituent of ER-mitochondria protein bridges, downregulation of phosphofurin acid cluster sorting protein-2 (*PACS2*) or mitofusin-2 (*MFN2*) has been shown to reduce MAMs (37–39). Mitochondria in CHLA15-shPACS2 cells with Pacs2 expression reduced >50% were larger than CHLA15-shCtrl cells but similar in roundness and circularity (Figure 5B-D). There was a trend toward reduced MAM per mitochondria though the distribution of MAMs to mitochondria was preserved (Figure 5F; Table 2). MAM length and proximity were reduced, with Pacs2 knock-down cells having a reduced MAM per mitochondrial perimeter across all gap widths (Figure 5F-G; Table 2). The proportion of MAM that came within 10nm or 25 nm of the OMM was reduced (p=0.02 and p<0.05, respectively), as was the proportion of the OMM perimeter in contact with a MAM at all gap-widths (Figure 5G). By comparison, mitochondria in CHLA15-shMFN2 cells with Mfn2 reduced ∼60% had reduced circularity and roundness (p=0.03 and p<0.01, respectively) but size was maintained (Figure 5B-E). There were fewer MAMs per mitochondria (p<0.01), and the frequency of mitochondria with ≥2 MAMs was reduced while those absent MAMs were increased (both p<0.01; Figure 5F). Moreover, MAM lengths were reduced across all gap-widths (Figure 5G). We tested the sensitivity of these cells to the Bcl2/Bclx-inhibitor, ABT-737, a pharmacological enhancer of MOMP in CHLA15 cells, and knock-down of Mfn2 or Pacs2 phenocopied the ABT-737 resistance seen in CHLA20 cells, shifting the IC_50_ >18-fold and 8-fold, respectively (Figure 5H). In addition, CHLA15-shMFN2 cells, which had more substantial MAM alterations than did CHLA15-shPACS2 cells, showed a resistance to tBid and BimBH3 in mitochondrial profiles that phenocopied multidrug resistant neuroblastomas (Figure 5I).

**Figure 5.**
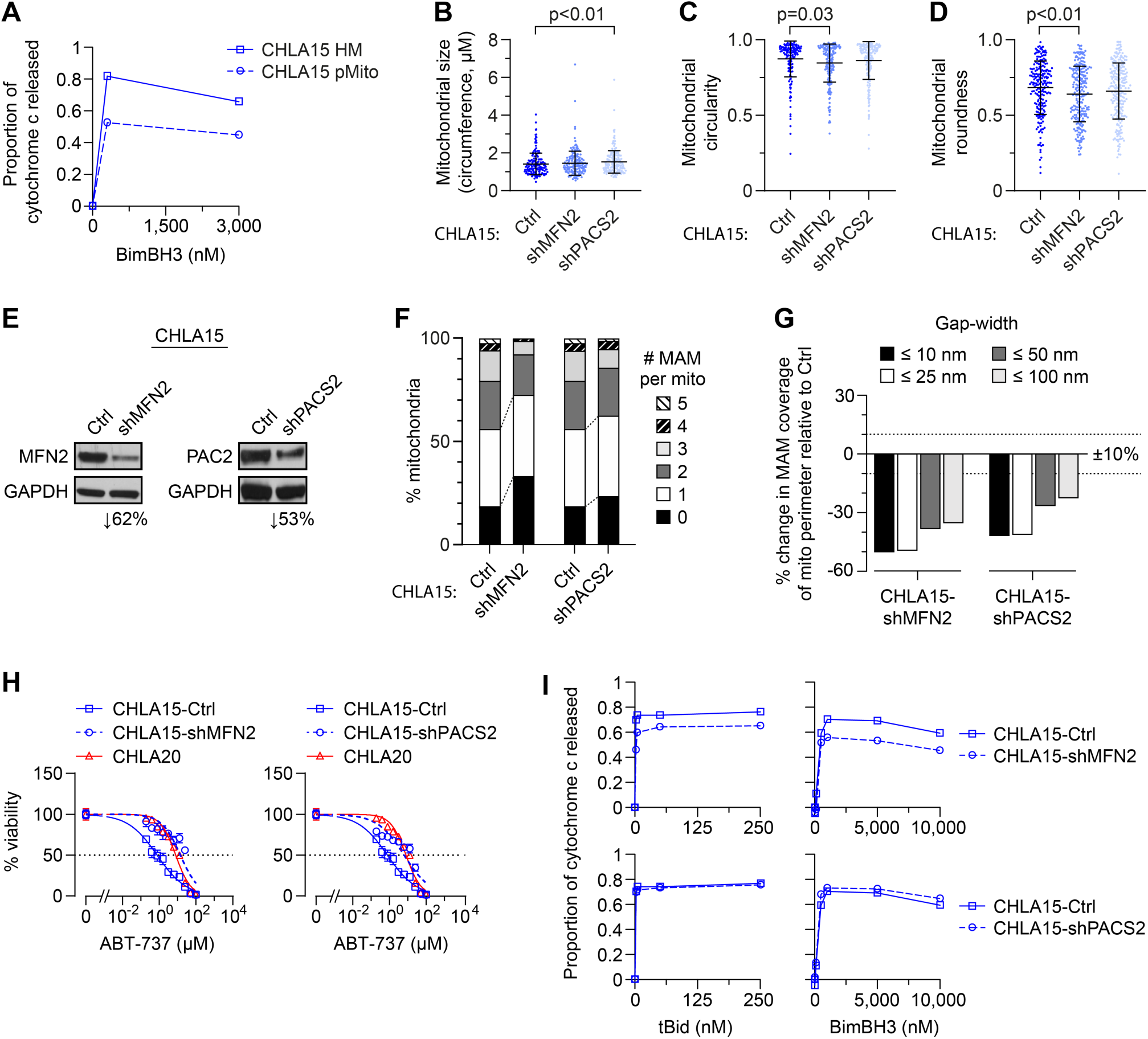
Chemically or genetically reducing MAMs leads to MOMP resistance. **(A)** Heavy membrane (HM) fractions of CHLA15 were rendered MAM-depleted by partial proteolysis and immune-magnetic cell sorting to derive purified mitochondria (pMito), and cytochrome c release measured in response to BimBH3 peptide. TEM analyses were used to quantify mitochondrial size **(B)**, circularity **(C)** and roundness **(D)** in CHLA15 cells transfected with a sh-control (Ctrl), shMFN2 or shPACS2 constructs (mean +/- SD shown); with immunoblot assessment of protein knockdown, **(E)**. Proportion of mitochondria with 0 to 5 MAMs are shown for each **(F)**; as is the percentage of the mitochondria perimeter with an apposed ER within defined gap-widths (as shown), **(G)**. For B-D and F-G, n=214-246 mitochondria per cell line. **(H)** In vitro viability of CHLA15-ctrl, CHLA15-shMFN2, CHLA15-shPACS2, and CHLA20 cells following 72hr exposure to the Bcl2/Bclx-inhibitor, ABT-737. **(I)** Mitochondrial cytochrome c release in response to tBid and BimBH3 peptide in CHLA15-Ctrl, CHLA15-shMFN2 and CHLA15-shPACS2 cells. For A and I, data points are mean of duplicate wells (SD<0.05 at all points) in a representative experiment from at least two biological replicates; for H, data points are mean and SD from triplicate wells, experiments are representative of at least two biological replicates. Statistical analyses were performed using an unpaired two-sided Student’s t-test, with significance p<0.05.

**Table 2.** Summary of CHLA15-Ctrl (DX) and CHLA15-shMFN2 and -shPACS2 models.

|  | **CHLA15-**<br>**ctrl** | **CHLA15-**<br>**shMFN2** |  | **CHLA15-**<br>**shPACS2** |  |
| --- | --- | --- | --- | --- | --- |
| **<u>MITOCHONDRIA:</u>** |  |  |  |  |  |
| Number analyzed | 214 | 234 |  | 246 |  |
| Perimeter, mean (nm) | 1,416 | 1,468 | *p*= ns | 1,530 | *p*<0.01 |
| Perimeter, median (nm) | 1,319 | 1,381 |  | 1,453 |  |
| Circularity, median | 0.918 | 0.892 | *p*=0.03 | 0.914 | *p*= ns |
| Roundness, median | 0.715 | 0.655 | *p*<0.01 | 0.686 | *p*= ns |
| % with 0 MAMs | 19% | 33% | *p*<0.01 | 24% | *p*= ns |
| % with 1 MAMs | 37% | 39% |  | 39% |  |
| % with 2-3 MAMs | 37% | 26% | *p*<0.01 | 22% | *p*= ns |
| % with ≥4 MAMs | 6% | 1% |  | 5% |  |
| %OMM with MAM gap-width ≤10nm | 1.7% | 0.9% | (↓47%) | 1.0% | (↓41%) |
| %OMM with MAM gap-width ≤25nm | 5.9% | 3.0% | (↓49%) | 3.5% | (↓41%) |
| %OMM with MAM gap-width ≤50nm | 12.6% | 7.8% | (↓38%) | 9.3% | (↓26%) |
| %OMM with MAM gap-width ≤100nm | 27.4% | 17.8% | (↓35%) | 21.2% | (↓23%) |
| **<u>MAM:</u>** |  |  |  |  |  |
| Number analyzed | 330 | 241 |  | 331 |  |
| MAMs per mitochondria | 1.54 | 1.03 | *p*<0.01 | 1.35 | *p*=0.07 |
| MAM frequency (per nm perimeter) | 918 | 1,425 |  | 1,137 |  |
| %MAM with gap-width ≤10nm | 22% | 17% | *p*=0.09 | 15% | *p*=0.02 |
| %MAM with gap-width ≤25nm | 47% | 43% | *p*= ns | 40% | *p*<0.05 |
| %MAM with gap-width ≤50nm | 68% | 73% | *p*= ns | 71% | *p*= ns |
| %MAM with gap-width ≤100nm | 100% | 100% |  | 100% |  |

## DISCUSSION

Most cancer deaths result from progression of disease that is resistant to available therapies yet the principal contributors remain incompletely understood. The diversity of targets and stress signals engaged by cancer treatments, and the myriad adaptations cancer cells use to subvert them, pose fundamental challenges. One effort to classify therapy resistance identifies mechanisms that reactivate, bypass, or induce indifference to a drug-perturbed pathway (40). This model is most useful for resistance to pathway-selective drugs, in particular, inhibitors of oncogenic kinases. Here we identify a novel convergence-based mechanism of resistance arising from attenuation of stress-induced MOMP as a consequence of reduced MAMs. Effective cancer therapies activate stress signals that converge on mitochondria (23, 26, 41–44). Mitochondrial sensitization toward MOMP is regulated through the activities of MAMs that serve as essential contacts for the exchange of metabolites between ER and mitochondria. Since mitochondria serve as a terminal node integrating stress signals to induce the death of the cancer cell via MOMP, their attenuated responsiveness contributes to multidrug resistance. Importantly, this mechanism is not exclusive to other contributors to therapy resistance operative upstream of mitochondria.

We identified the MOMP-attenuated phenotype measuring mitochondrial responses to the Bak and/or Bax activators, tBid and Bim. Mitochondrial profiling was developed by Letai (23) and has been used to define Bcl2-family dependencies and predict responses to chemotherapy (42–44) and molecularly targeted drugs (42, 45). We optimized this for the study of neuroblastoma, a tumor that often responds to chemoradiotherapy with tumor regression (46) followed by lethal relapse (12). Limited availability of paired clinical samples has constrained such investigations, so we derived patient-matched tumor models at both diagnosis and relapse, the latter frequently manifesting multidrug resistance (10). Cell lines were used as they can be established under identical conditions from tumor biopsies, or from tumor-involved bone marrow, the latter the most common source at relapse. We show that mitochondria from relapsed neuroblastomas have markedly reduced MOMP responses to tBid and Bim (14) compared with their at-diagnosis counterparts, providing evidence that attenuated mitochondrial signaling is acquired under the selective pressure of multimodal therapy. Attenuated MOMP was reproducibly identified in nearly all relapsed neuroblastomas in response to either death-activator, although tBid was more potent than BimBH3. Indeed, BimBH3 is an intrinsically unstructured peptide (47) that typically engages Bak or Bax at micromolar exposures, while recombinant tBid protein binds lipid membranes to activate Bak or Bax at nanomolar exposures, highlighting the important role of lipid membranes in facilitating MOMP (15, 48). Quantitative variability among biological replicates were notable, reflecting myriad events impinging on Bak/Bax-mediated pore formation, including temperature, p*H*, detergents and metabolites (49). Working with patient-matched pairs in parallel controlled for many of these variables and relative differences between tumors were robust.

Our work utilized neuroblastoma cell lines that represent oligoclonal outgrowths from explanted tumors. If MOMP sensitivity were highly heterogeneous within a tumor, we would predict some diagnosis-relapse pairs to have attenuated mitochondrial responses present at diagnosis. Instead, our data support that contributions from intra-tumoral heterogeneity are minor in comparison with the effect of therapeutic selective pressure. We confirmed that attenuated mitochondrial responses correlated with resistance to chemotherapy and ionizing radiation. For the latter, differential cell death follows equivalent genotoxic stress supporting a post-target resistance mechanism. Resistance is also conferred to molecularly targeted drugs such as Bcl2 and Alk inhibitors, even in the absence of prior exposure, since these operate upstream of mitochondria. More surprising, however, is evidence that selecting for resistance to such target-selective agents in vitro can induce the attenuated MOMP phenotype accompanied by multidrug resistance. This provides a potential mechanism for the significant proportion of patients with emergent resistance to therapeutic kinase inhibitors that have no on-pathway resistance mechanism identified (3–6, 8), a clinically relevant finding. That mitochondrial responses can discriminate chemosensitive from chemoresistant tumors has been shown using similar approaches (43), which also demonstrate a mitochondrial basis for the selective killing of tumor cells by chemotherapy (26, 44). Here, we expand this notion to demonstrate a mitochondrial basis for multidrug resistance in response to therapeutic pressure during multimodal therapy.

We identified reductions in MAM content from tumors with attenuated MOMP when visualized by fractionation or quantified in electron micrographs. ER-mitochondria contacts at MAMs regulate fission/fusion dynamics to optimize mitochondrial shape (50), which impacts Bax-induced MOMP in murine hepatocytes and murine embryo fibroblasts (MEFs) by altering cooperation among Bcl2-family members. Smaller mitochondria with increased membrane curvature are more resistant to stress-induced MOMP (51), and are capable of promoting incomplete-MOMP followed by mitochondrial re-population and cell survival following stress (52), possibly contributing to chemoresistance (51). Consistent with this, the CHLA15/CHLA20 pair had smaller more circular mitochondria in the post-relapse MOMP-resistant tumor, however, this was absent in SKNBE1/SKNBE2C (in which mitochondria in the relapsed model were larger). While mitochondrial size, shape, mass and mtDNA content differed variably, only the reduction in MAM content and ERMCs was consistently present across models with attenuated mitochondrial responses, while absent in the sole pair without this feature.

Our image analyses were done in two dimensions yet inter-organelle contacts occur in 3D, so to ensure representative findings we quantitatively analyzed >2,400 MAMs from >1,700 mitochondria across 9 tumor models. In both the CHLA15/CHLA20 and SKNBE1/SKNBE2C pairs, the ER-mitochondrial connectedness was markedly reduced in the therapy resistant post-relapse model, with significantly more orphan mitochondria and fewer mitochondria with ≥2 MAMs. Moreover, the proportion of mitochondrial surface with an apposed MAM was reduced. In contrast, the only tumor pair without attenuated MOMP responses or chemoresistance, CHLA122/CHLA136, did not have such alterations. That MAMs might be reduced in cancer cells was originally posited by Howatson and Ham from EM studies of mouse liver tumors (53). An essential role for MAMs in mediating apoptotic sensitivity has been demonstrated by Chipuk et al (30). Using MEF heavy-membrane mitochondria-enriched fractions, they showed that MOMP sensitivity to Bid peptides was attenuated when mitochondria were purified from their associated MAMs, and rescued by the add-back of MAM-containing heterotypic membranes (30). Consistent with this, we could phenocopy attenuated MOMP responses by reducing MAMs from mitochondria in diagnostic-tumors using chemical proteolysis or genetic perturbation. In the latter, we confirmed that a MAM-reduced phenotype was achieved using EM image analysis.

Collectively, these data support an essential factor for mitochondrial sensitization to MOMP is provided by MAMs. Korsmeyer’s group showed that many intrinsic cell death signals require both functional Bak or Bax *and* ER-released Ca^2+^ (54) and deficiencies in MAM-derived Ca^2+^ transfer can confer apoptosis resistance in mesothelioma cells (55). We found a >3-fold increase in Ca^2+^ transfer coupling time in therapy resistant SKNBE2C (REL) cells compared with DX cells, which was normalized using a genetic linker to tighten MAM contacts. However, we found no change in coupling time for the CHLA15/CHLA20 pair. Our EM analyses provide a potential explanation as CHLA20 has MAM reductions more limited to tighter gap-widths with relative preservation at larger gap-widths capable of creating Ca^2+^ transfer microdomains (34, 56). While altered Ca^2+^ transfer may contribute to MOMP regulation, it is not indispensable for the attenuated MOMP sensitivity in relapsed neuroblastoma. While narrow gap-width MAMs are unable to accommodate the Ca^2+^ transfer machinery, such close contacts are critical for phospholipid transport. MAMs are enriched in lipid synthesizing enzymes, mitochondria are dependent on ER-derived lipid transport, and the composition of mitochondrial membranes is a determinant of MOMP sensitivity. Chipuk and Green demonstrated that the reduced MOMP sensitivity of mitochondria isolated from MAMs can be restored by the return of MAMs themselves, or by the provision of sphingosine-1-phosphate or hexadecenal, which support Bak or Bax oligomerization (30). Indeed, such tight contacts (<10 nm) have been referred to as lipid-ERMCs, reflecting this essential role (57, 58), and reductions have been associated with altered mitochondrial lipid composition in neurons in Alzheimer models (57). It is intriguing to speculate that reduced MAMs contribute to stress resistance in cancer cells by altering the mitochondrial OMM sphingolipid environment to attenuate Bak and Bax responsiveness, and further investigation is warranted (59).

We propose that MAMs serve as physiological regulators of apoptosis in cancer cells and that MAM reductions comprise an adaptive response to therapeutic stress, providing a novel mechanism for multidrug resistance involving disrupted ER-mitochondria communication (Figure 6). During cancer therapy, cells with reduced MAMs have a higher survival probability via “incomplete-MOMP” (52) leading to selective enrichment. Alterations in MAMs have increasingly been linked to diverse pathological states, including cardiomyocyte recovery after re-perfusion (60), ototoxic injury (61), neurodegeneration (20-22, 62, 63), and obesity-related diabetes (19, 64). Moreover, our attenuated MOMP model proposes that multidrug resistance does not reflect an absence of activated death signals but their insufficiency to trigger MOMP. Investigational cancer drugs are first tested in patients with multidrug resistance following failed therapies. For the many drugs that do not induce a tumor response in such studies, we cannot differentiate those that effectively engaged their target and liberated death-activating signals (that remained subthreshold) versus those that did not, leading investigators to abandon some drugs that might have clinical efficacy in other settings. Importantly, recognizing this phenotype can facilitate the development of tools to measure ER-mitochondria interactions for clinical use in predicting therapy resistance, and provides a novel framework for testing interventions to prevent emergent resistance or restore mitochondrial competence to resistant cancers.

**Figure 6.**
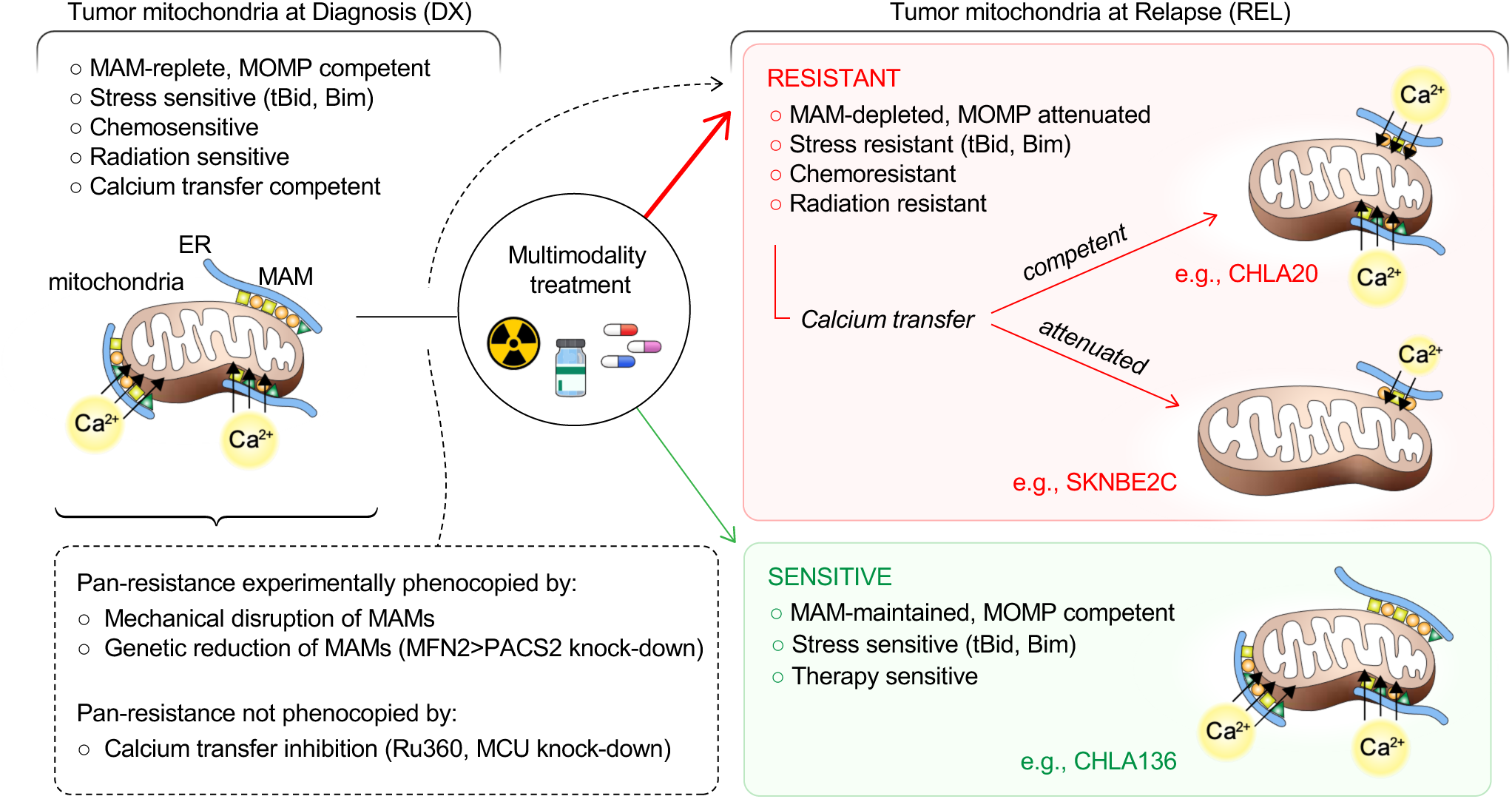
MAM-mediated mitochondrial resistance model. At diagnosis (DX), neuroblastoma cells have MAM-replete mitochondria that are MOMP competent, and relatively sensitive to chemotherapy, radiation therapy, and targeted agents. Relapsed (REL) tumor cells recurring after multimodal therapy have reduced MAMs (by number and/or proximity to mitochondria) and are relatively multidrug resistant as a consequence of mitochondria that have attenuated MOMP in response to stress signals (red box). Rare tumors at relapse retain MOMP competence and therapy sensitivity (green box). Calcium transfer from ER to mitochondria may be reduced when MAMs are markedly attenuated but this reduced transfer is not essential for the multidrug resistant phenotype. Transfer of additional metabolites like phospholipids may also contribute. MOMP, mitochondrial outer membrane permeability.

## METHODS

### Cell Lines

Neuroblastoma cell lines were obtained from the COG/ALSF Childhood Cancer Repository (www.CCcells.org). Cells were grown in IMDM Media 12440 (Life Technologies, Gaithersburg, MD) supplemented with 10% fetal bovine serum, 2mM L-glutamine, 1% ITS, 100 U/ml of penicillin and 100 mcg/ml gentamycin. Cell lines obtained at diagnosis (DX) were kept physically separate from those at relapse (REL) at all times to avoid cross-contamination. Cell lines are referred to by name, often with the descriptor “DX” or “REL” to note their time of derivation. All cell lines were subjected to STR identity and pathogen testing every 4-6 months. Tissue culture conditions were 37°C in a humidified atmosphere of 5% CO_2_. Cell lines were interrogated for cancer gene mutations using the FoundationOne CDx assay (Foundation Medicine) to confirm *TP53*, *ALK* and additional cancer gene mutation status, including variant allele frequencies.

### Isolation of mitochondria-enriched heavy membrane fractions

Heavy membrane fractions were obtained by rupturing cells resuspended in isolation buffer A (250mM Sucrose, 20mM HEPES, 1mM DTT, 10mM KCl, 1mM EDTA, 1mM EGTA, 1.5mM MgCl_2_) with protease inhibitor cocktail by 20 strokes in a 2 mL glass Dounce homogenizer, followed by removal of cell debris and nuclei by centrifugation of supernatants at 800g for 10 min and then 1,050g for 10 min at 4°C. Mitochondria-enriched fractions were collected by centrifuging the supernatant of the second spin at 12,000g for 10 mins at 4°C.

### Magnetic Activated Cell Sorting (MACS) to purify mitochondria

To physically separate MAMs from crude mitochondria, we used MACS according to manufacturer’s MidiMACS^TM^ Starting Kit (LS) protocol (130-042-301, Mitenyi Biotec, Germany). Cells were collected in cold PBS and re-suspended in cold Buffer A with protease inhibitor at the final concentration of 10 million cells/ml. After 20-25 strokes of homogenization using a 7ml Dounce homogenizer, 1ml of cell lysate was mixed with 9ml of ice-cold 1X Separation Buffer (130-091-221, Mitenyi Biotec) and incubated with 50μl anti-TOM22 microbeads at 4°C on a shaker for 1 hour. The mixture was passed stepwise through an LS column (130-042-401, Mitenyi Biotec), pre-rinsed with 3ml of 1X Separation Buffer and installed in the magnetic field of the MidiMACS Separator (130-042-302, Mitenyi Biotec). After washing three times with 3ml of 1X Separation Buffer, the LS column was removed, and magnetically labeled mitochondria flushed into a collection tube using 1.5ml of 1X Separation Buffer. Purified mitochondria were used immediately for analysis or centrifuged at 13,000g for 2 minutes at 4°C and stored in 100μl storage buffer on ice.

### Mitochondrial profiling

Heavy membrane or purified mitochondria cell fractions were resuspended in buffer C [10 mM Tris-MOPS (pH 7.4), 1 mM KH2PO4, 10 mcM Tris-EGTA, 5 mM glutamate, 2 mM malate, 125 mM KCl] at a final concentration of 1μg/μl. For cytochrome c release reactions, 50μl of mitochondria (1μg/μl protein) were treated with Caspase-8-cleaved tBid from 5-500nM (882-B8-050, R&D Systems, Minneapolis, MN), Bim-BH3 from 10-10,000nM (Ac-DMRPEIWIAQELRRIGDEFNAYYARR-amide; New England Peptide, Gardner, MA) or 300μM BidAltBH3 (Ac-EDIIRNIARHAAQVGASMDR-amide; New England Peptide, Inc.). Reactions were incubated at 37°C for 30 mins and spun at 12,000g for 10 min at 4°C: 10μl of the supernatant and the 10 μl pellet (mitochondrial) fractions were resuspended in 50μl of 0.1% Triton X in PBS in duplicate in Quantikine, Human Cyto C Immunoassay 96-well plates followed by ELISA detection (R&D Systems, Minneapolis, MN). The fractional release of mitochondrial cytochrome c was calculated for each condition as [mean intensity_supernatant_/(mean intensity_mitochondria_ + mean intensity_supernatant_)]. We report % cytochrome c release normalized to the non-specific release induced by treatment with DMSO (control) as [(fractional release_treatment_ -fractional release_dmso_)/(1-fractional release_dmso_)]*100. Experiments with <30% cytochrome c release to DMSO and BidAltBH3 peptide were analyzed.

### MAM/mitochondria separation

Heavy membrane fractions were resuspended in 2ml Isolation Medium (250 mM Mannitol, 5 mM HEPES pH 7.4, 0.5 mM EGTA, 0.1% BSA) with freshly added protease inhibitors and slowly layered onto 8ml of 30% Percoll gradient prepared with Gradient Buffer (225 mM Mannitol, 25 mM HEPES pH 7.5, 1 mM EGTA, 0.1% BSA) in an Ultra-Clear Beckman Centrifuge Tube, and spun at 95,000g for 60 min to resolve MAM from more pure mitochondria, as described (29).

### Cytotoxicity assays

Cells were seeded into Corning® 96-well Flat Clear Bottom White Polystyrene TC-treated luminescent microplates (3610, Corning, Corning, NY) in triplicate at a density of 10,000 cells/well and allowed a 24 hour recovery period. Cells were treated with 100ul of control Gibco^TM^ IMDM full culture media (12440-053, Gibco by Life Technologies, Carlsbad, CA). ABT-737 was tested from 1nM-200µM, added to IMDM culture media for 48 hours. Chemotherapy exposure was 72 hours, concentrations tested were: etoposide (341205-25MG, EMD Millipore, Billerica, MA) 10nM-10µM, cisplatin (63323-103-51, Fresenius Kabi, Lake Zurich, IL) 0.5nM-75µM, carboplatin (216100-25MG, EMD Millipore) 6nM-10µM, doxorubicin (5927S, Cell Signaling Technology, Danvers, MA) 0.5nM-10µM, and masfosfamide (M110300 Toronto Research Chemicals, Inc, Toronto, Canada) 2nM-75 µM. ALK inhibitors [crizotinib (C-7900, LC Laboratories, Woburn, MA), ceritinib (C-2086, LC Laboratories), and lorlatinib (5640, Tocris Bioscience, Minneapolis, MN)] were tested from 0.01nM-10µM and assessed after 5 days. Cells were irradiated on a Cs-137 Gammacell 40 irradiator S/N 186 (Nordion Ltd, Kanata, Ontario, Canada) at a dose rate of 1.3 cGy/sec from 0.5 to 5 Gy and assessed after 7 days. Viability was assessed using CellTiter-Glo® Assay according to the manufacturer’s instructions (Promega, Madison, WI; #G7571). A Synergy™2 microplate reader with Gen5™ software (BioTek Instruments, Winooski, VT) was used to measure luminescence. For each plate, mean relative luminescence was calculated from at least three technical replicates, normalized to control samples. Nonlinear regression algorithms in Prism software (GraphPad8) were used to calculate IC_50_ values.

### Real-time qPCR detection of mtDNA

Mitochondrial DNA content was quantified by qPCR to define the mtDNA/nucDNA ratio using each of two mitochondrial-genome (MT-C01 and MT-ATPase6) and two nuclear-genome (CFAP410 and MTTP) genes. Assays were run in triplicate using Taqman Gene Expression Mastermix (4369016, Applied Biosystems) and the following primer/probes from Invitrogen: MT-CO1 (Hs02596864_g1), MT-ATPase6 (Hs02596862_g1), CFAP410 (Hs00223770_cn), and MTTP (HS0405900_cn). mtDNA/nucDNA ratio was determined as the mean and SD of all biological replicates.

### Citrate Synthase measurements

Mitochondrial mass was assessed by measuring citrate synthase activity from 4 million whole cells normalized to protein input (U/mg) in triplicate from three biological replicates using the Citrate Synthase Assay Kit (CS0720, Sigma-Aldrich, St, Louis, MO).

### Electron microscopy

Intact cells for electron microscopic examination were fixed with 2% glutaraldehyde with 1% tannic acid in 0.1M sodium cacodylate buffer, pH 7.4, overnight at 4°C. After buffer washes, the samples were post-fixed in 2.0% osmium tetroxide with 1.5% K_3_Fe(CN)_6_ for 1 hour at room temperature, and rinsed in ddH_2_O prior to en bloc staining with 2% uranyl acetate. After dehydration through a graded ethanol series, the tissue was infiltrated and embedded in EMbed-812 (Electron Microscopy Sciences, Fort Washington, PA). Thin 70 nm sections were stained with uranyl acetate and SATO lead and examined with a JEOL 1010 electron microscope (JEOL, Peabody, MA) at 70 kV fitted with a Hamamatsu digital camera and AMT Advantage NanoSprint500 software.

### Measurement of MAMs, ER-mitochondria gap distances, and mitochondria circularity and roundness

TEM images at 50,000x magnification were analyzed blinded in Image J by hand-masking the mitochondria perimeter and ER interface. Analysis of the ER-mitochondria interfaces were extracted with a custom macro (65) available at: http://sites.imagej.net/MitoCare/ that quantifies ER interface metrics binned within a 10, 25, 50 or 100 nm gap distance from the mitochondria (34).

### Detection of ER-derived calcium transfer into mitochondria

Mitochondria-targeted calcium construct 4mtGCamp6f was electroporated into cells with with Amaxa Nucleofactor (Lonza, Basel, Switzerland). Cells were allowed to recover in full IMDM media for 24 hours and then plated on poly-D lysine covered slides, washed three times with Ca2+-free HBSS medium (Life Technologies Inc., Grand Island, New York) and loaded with 1μM Fura 2-AM (Life Technologies Inc., Eugene, OR) cytoplasmic calcium indicator in the presence of 0.03% Pluronic F127 and 100μM sulfinpyrazone at 35 °C for 30 min. Fura-2AM-loaded cells were washed with Ca^2+^-free HBSS medium and mounted into microscope loading chambers. Measurements of [Ca^2+^]_m_ and [Ca^2+^]_c_ were carried out using a multiwavelength beamsplitter/emission filter combination and a high quantum-efficiency cooled CCD camera. Excitation at 340 nm was used for Fura-2AM and, 380 nm was used for 4mtGCamp6f. Basal fluorescent levels were measured for 120 ms, after which calcium release from ER was evoked by addition of 100μM carbachol. To determine the average [Ca^2+^]_c_ and [Ca^2+]^ signals, mean fluorescence intensities (ratios of Fura-2AM to 4mtGCamp6f) were calculated for essentially all individual whole-cell areas in each run after subtraction of the background fluorescence measured at cell-free areas of the field. The *K*d values of 340nm for Fura-2 AM and 380nm for GCamp6f at 50% maximal release were used to calculate the coupling time.

### Plasmids, retroviral constructs, and reagents

A synthetic mitochondria-ER linker was used to physically recouple ER and mitochondria, as described (34). Lentiviral shRNAs to MFN2 (RHS5086-EG9927-GIPZ MFN2), PACS2 (RHS5086-EG23241-GIPZ PACS2) and GIPZ non-silencing lentiviral shRNA Control were purchased from GE Health Dharmacon, Inc., Lafayette, CO. 4mtGCamp6f was provided by Dr. Diego De Stefani. Primary antibodies: anti-BAP31 (ab109304, Abcam Inc, Cambridge, MA), anti-β-tubulin (T8328, Sigma-Aldrich, St. Louis, MO), anti-calnexin (C7617, Sigma-Aldrich), anti-FACL4 (22401-1-AP, Proteintech, Rosemond, IL), anti-GAPDH (2118, Cell Signaling Technologies, Danvers, MA), anti-*γ*-H2AX (NB100-384, Novus, Littleton, CO), anti-MFN2 (ab56889, Abcam Inc), anti-MCU (HPA016480, Sigma-Aldrich), anti-noxa (OP180, Calbiochem/EMD Chemicals, San Diego, CA), anti-PACS2 (ab129402, Abcam Inc), anti-p53 (sc-126, Santa Cruz Biotechnology, Dallas, TX), anti-p21 (sc-6246, Santa Cruz Biotechnology), anti-TMX1 (256-270, SAB1105403, Sigma-Aldrich), and anti-TOMM40 (18409-1-AP, Proteintech, Rosemond, IL). To quantify protein knockdown, densitometry was performed using ImageJ. The intensity of each band was normalized to its respective loading control for comparison.

### DNA damage detection

10,000 cells were plated onto poly-L lysine coated microscope slides, allowed to adhere 24 hours and irradiated at 2Gy for 1 hour on a Cs-137 Gammacell 40 irradiator S/N 186 (Nordion Ltd, Kanata, Ontario, Canada) at a dose rate of 1.3 cGy/sec. Cells were washed once with warm PBS and fixed with pre-warmed paraformaldehyde for 10 minutes, permeabilized with 2% Triton-PBS for 5 minutes at 4°C, washed twice with PBS-5% triton and blocked for 10 minutes in Duolink blocking solution. Cells were then incubated for 1 hour at 37°C with *γ*-H2AX primary antibody (NB100-384, Novus Biologicals, Centennial, CO) at 1:1000 dilution, washed twice with PBS-T and incubated at 37°C with AlexaFluor goat anti-rabbit 488 (ab150077; Abcam Inc, Cambridge, MA) at 1:500 dilution for at 45 minutes. After washing 4 times with PBS-T they were stained with VectaShield DAPI staining, covered with a microscope coverslip, and analyzed using a Leica DMR fluorescent microscope at 40X magnification and the number of *γ*-H2AX foci/cell quantified.

## Statistical analyses

Statistical comparisons for mitochondrial size, roundness, circularity and MAM content per mitochondria were performed with the Mann-Whitney U test, 2-tailed; comparisons for cytochrome c release, DNA damage foci, Ca^2+^-transfer coupling time, mtDNA content and mass were performed with the student t-test, independent values, two-tailed; comparisons for the proportion of mitochondria with diverse tether numbers were performed using the Chi-Square test. For all, significance was defined as p<0.05, trend p<0.10, and ns p≥0.10.

## AUTHOR CONTRIBUTIONS

JC, DMB, JV, CPR, JS, GH and MDH designed the experiments; CPR provided tumor models and model validation; JC, DMB, KCG, MCP, JV, KL, JCY, MC, ES, PD, ENO, ELC, JS and AV conducted the experiments and acquired data related to ER-mitochondria connectivity, signaling, and stress response; JC, DMB and GH conducted the experiments and acquired data related to calcium transfer; YL performed statistical analyses; JC, DMB, MCP, JV, KL, JCY, ECL, ES, AV, CPR, JS, GH and MDH analyzed the data; and JC, JS, DMB, CPR, KCG, GH and MDH wrote and edited the manuscript.

## ACKNOWLEDGEMENTS

This paper is dedicated to the memory of our dear friend and co-author Madison Pedrotty. We thank Renata Sano (Children’s Hospital of Philadelphia) for stimulating discussions related to this work, the Childhood Cancer Repository powered by Alex’s Lemonade Stand Foundation (www.cccells.org) for tumor model support, Diego De Stefani for provision of the GCamp6f plasmid, John Maris for cancer gene sequencing data, and the children and families providing tumor samples for research via the Children’s Oncology Group (COG). This study was supported by NIH CA198430, Alex’s Lemonade Stand Foundation, St. Baldrick’s Foundation, and CURE Childhood Cancer Foundation (to MDH); NIH CA216254 (to GH); and the Czech Science Foundation (No. GJ20-00987Y) to JS.

---

**Supplemental Figure 1.**
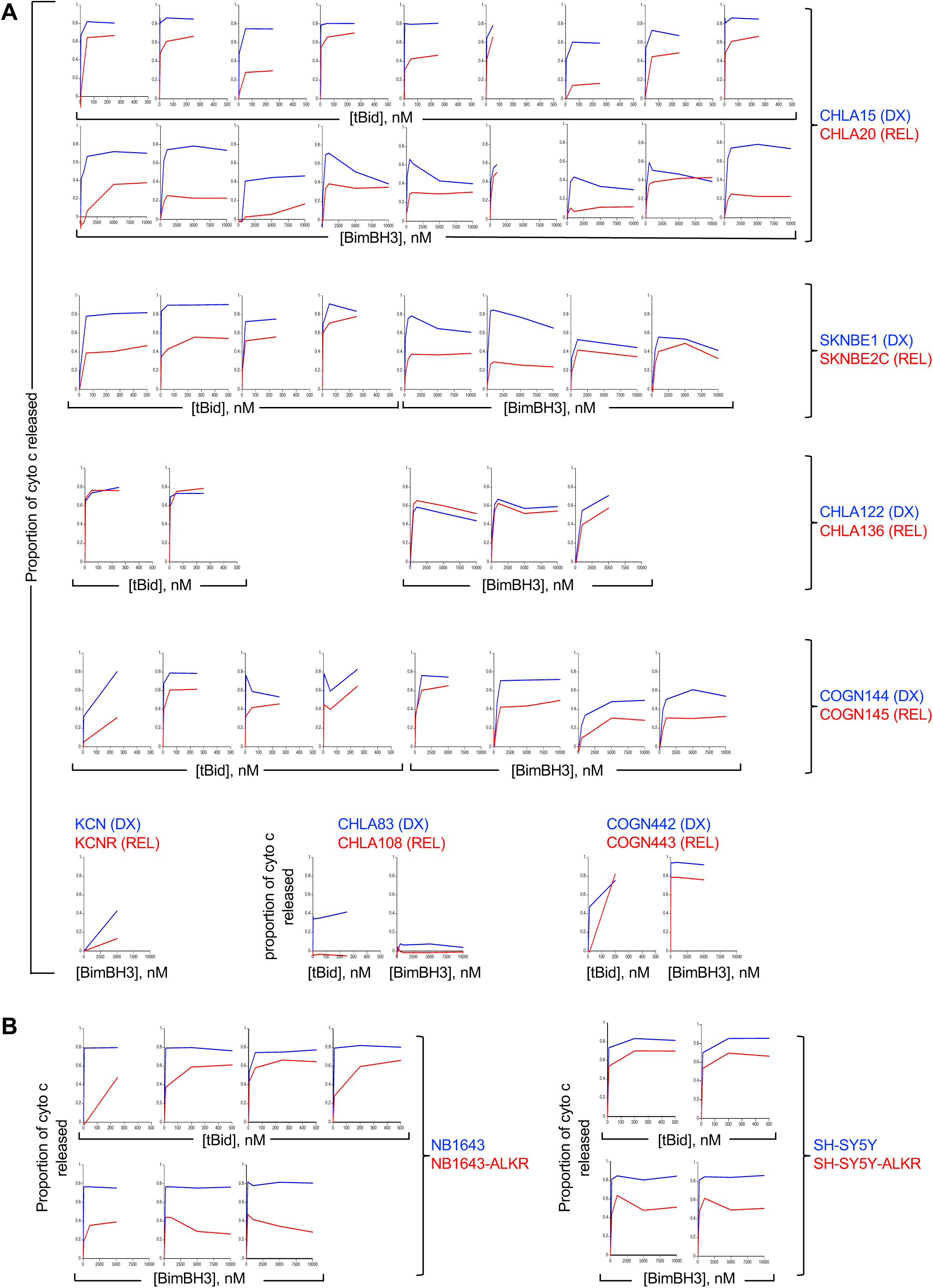
Mitochondrial cytochrome c release curves for neuroblastoma cell lines. The major therapy-induced stress signals, tBid and Bim (here recapitulated by the death-domain peptide itself, BimBH3), are capable of triggering MOMP in isolated tumor mitochondria through the activation of Bak and/or Bax. **(A)** Heavy-membrane mitochondria-enriched fractions were isolated at diagnosis (DX, blue curves), and from the same patient at relapse after failed multimodal therapy (REL, red curves). Model pairs that expanded readily and had spontaneous release of cytochrome c of <30% of that available (in response to control buffer or a death-domain inactivated BimBH3) were repeatedly assayed to demonstrate reproducibility, as shown. The CHLA122/CHLA136 pair did not demonstrate attenuated mitochondrial responses to tBid or BimBH3 in serial assays so was included in our mitochondrial characterization assays as a control, along with CHLA15/CHLA20 and SKNBE1/SKNBE2C as pairs manifesting profound MOMP attenuation. **(B)** A similarly attenuated cytochrome c release in response to tBid or BimBH3 was seen from heavy-membrane mitochondria-enriched fractions obtained from neuroblastoma cells selected for resistance to the Alk inhibitor, crizotinib (both SY5Y and NB1643 harbor activating ALK mutations sensitizing cells to crizotinib and other Alk inhibitors). Cells with crizotinib resistance are designated by “-ALKR” and are shown relative to parental crizotinib sensitive cells. Each plot represents a biological replicate experiment in which cytochrome c release represents the mean of 2 technical replicates (SD<0.05 at all points) and all biological replicates are shown.

**Supplemental Figure 2.**
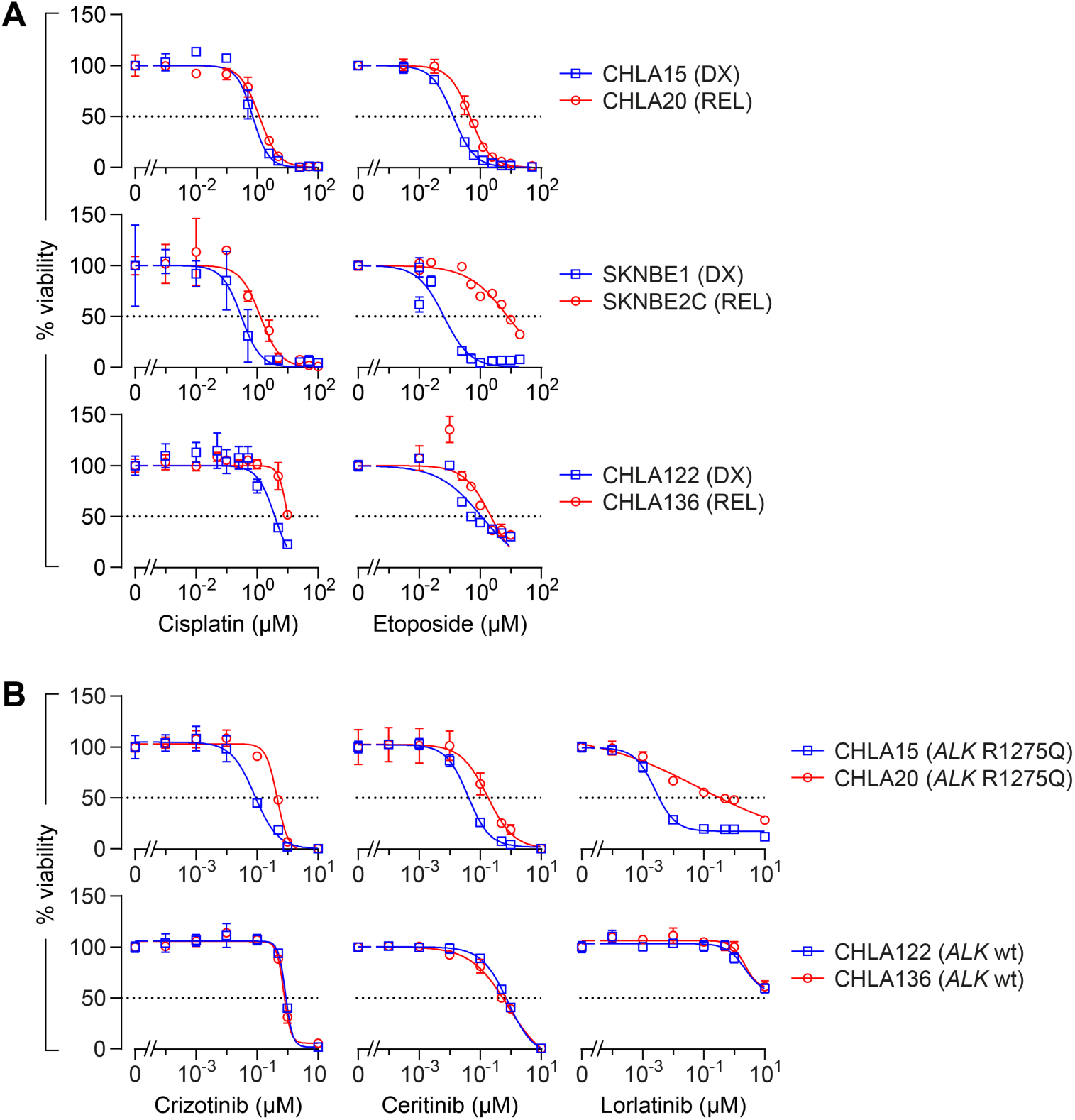
Most REL neuroblastomas are resistant to chemotherapeutics and molecularly targeted drugs. **(A)** In vitro viability for multiple DX/REL tumor pairs exposed to cisplatin or etoposide for 72 hours. **(B)** Viability is shown for DX/REL pairs with an activating ALK mutation (CHLA15/CHLA20) and without (CHLA122/CHLA136) following 120 hour exposure to the Alk inhibitors crizotinib, ceritinib or lorlatinib.

**Supplemental Figure 3.**
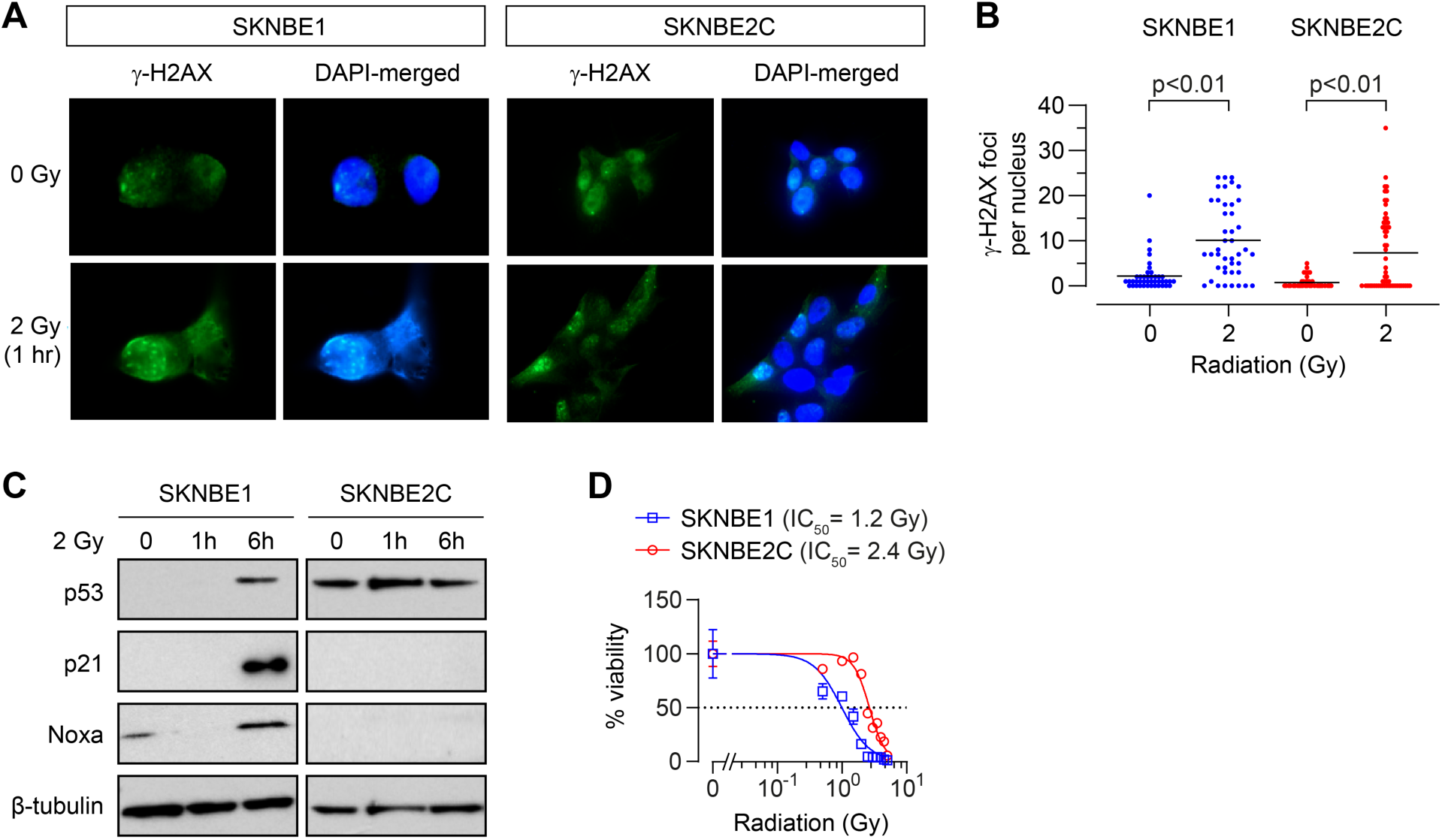
REL neuroblastoma cells are resistant to the cytotoxic effects of ionizing radiation despite incurring an equivalent increase in DNA damage. DNA-damage induced by 2 Gy ionizing radiation as measured by γ-H2AX foci at 1hr (n=42-58 nuclei/condition; mean shown) for SKNBE1 (DX, *TP53* wild-type) and SKNBE2C (REL, *TP53* mutant). Fluorescence detection of γ-H2AX foci and DAPI nuclear counterstain were assessed, with representative images shown **(A)**, and foci frequency plotted with mean for each condition, **(B**). Statistical analyses were performed using an unpaired two-sided Student’s t-test. **(C)** Cell lysates from cells treated with 2 Gy of ionizing irradiation were immunoblotted at 3 time-points for p53 and p53 target-genes with β-tubulin as a protein loading control. **(D)** In vitro viability following ionizing radiation at 7 days. For D: data points are mean and SD from triplicate wells, experiments are representative of at least three biological replicates.

**Supplemental Figure 4.**
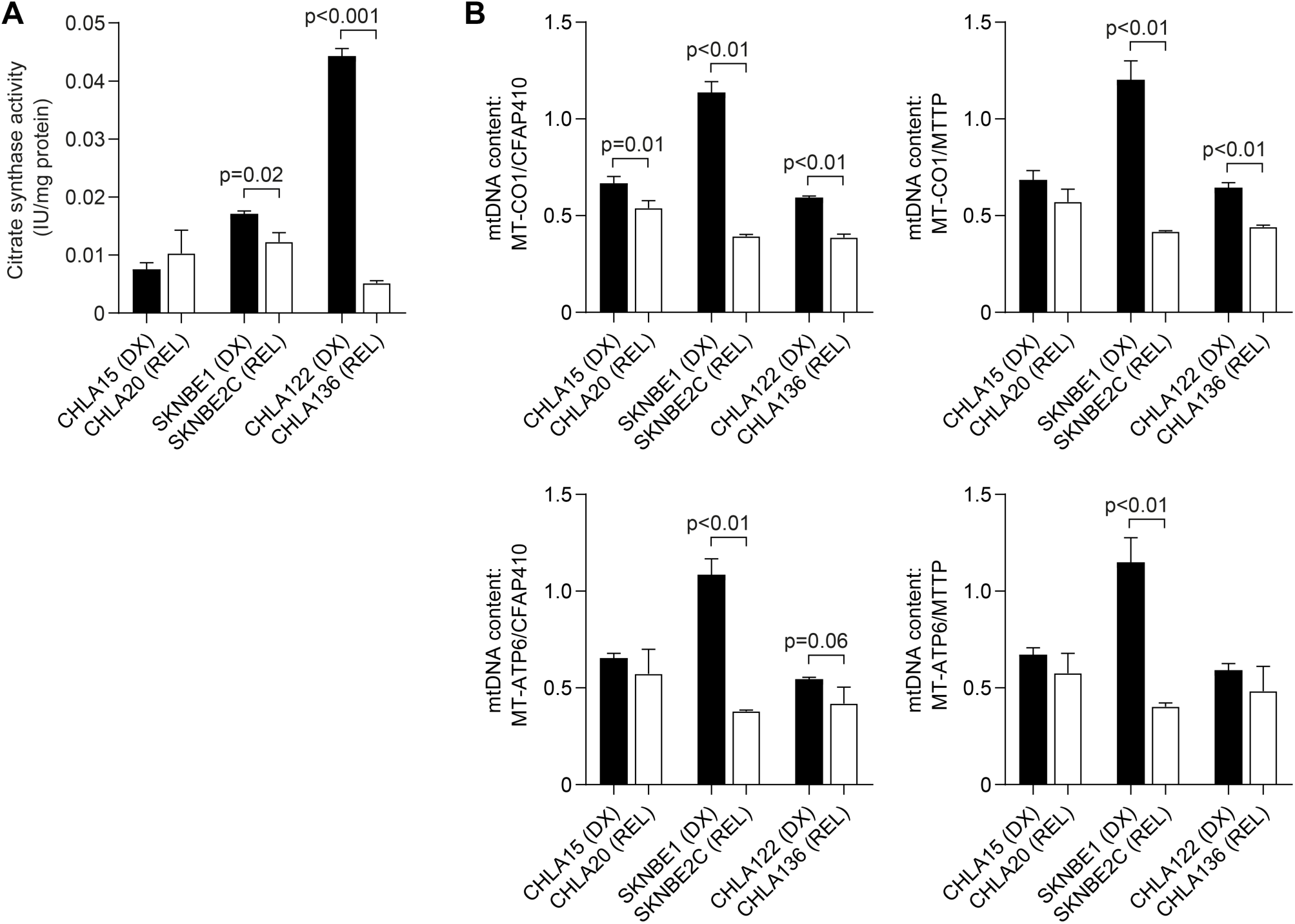
Neuroblastoma DX/REL pairs have altered mitochondrial biomass or morphology, but no consistent change is correlated with attenuated mitochondrial responses to stress, or chemotherapy resistance. **(A)** Mitochondrial biomass as assessed by citrate synthase activity across at-diagnosis (DX) and post-relapse (REL) pairs normalized to total protein input (IU/mg). **(B)** Mitochondrial DNA content as quantified by qPCR using the mtDNA/nucDNA ratio for each of two mitochondrial-genome (*MT-C01* and *MT-ATPase6*) and two nuclear-genome (*CFAP410* and *MTTP*) genes. The reduction in mtDNA in SKNBE2C is likely a consequence of mutant p53 signaling, as this has been reported in other cancer cell lines. Statistical analyses were performed using an unpaired two-sided Student’s t-test.

**Supplemental Figure 5.**
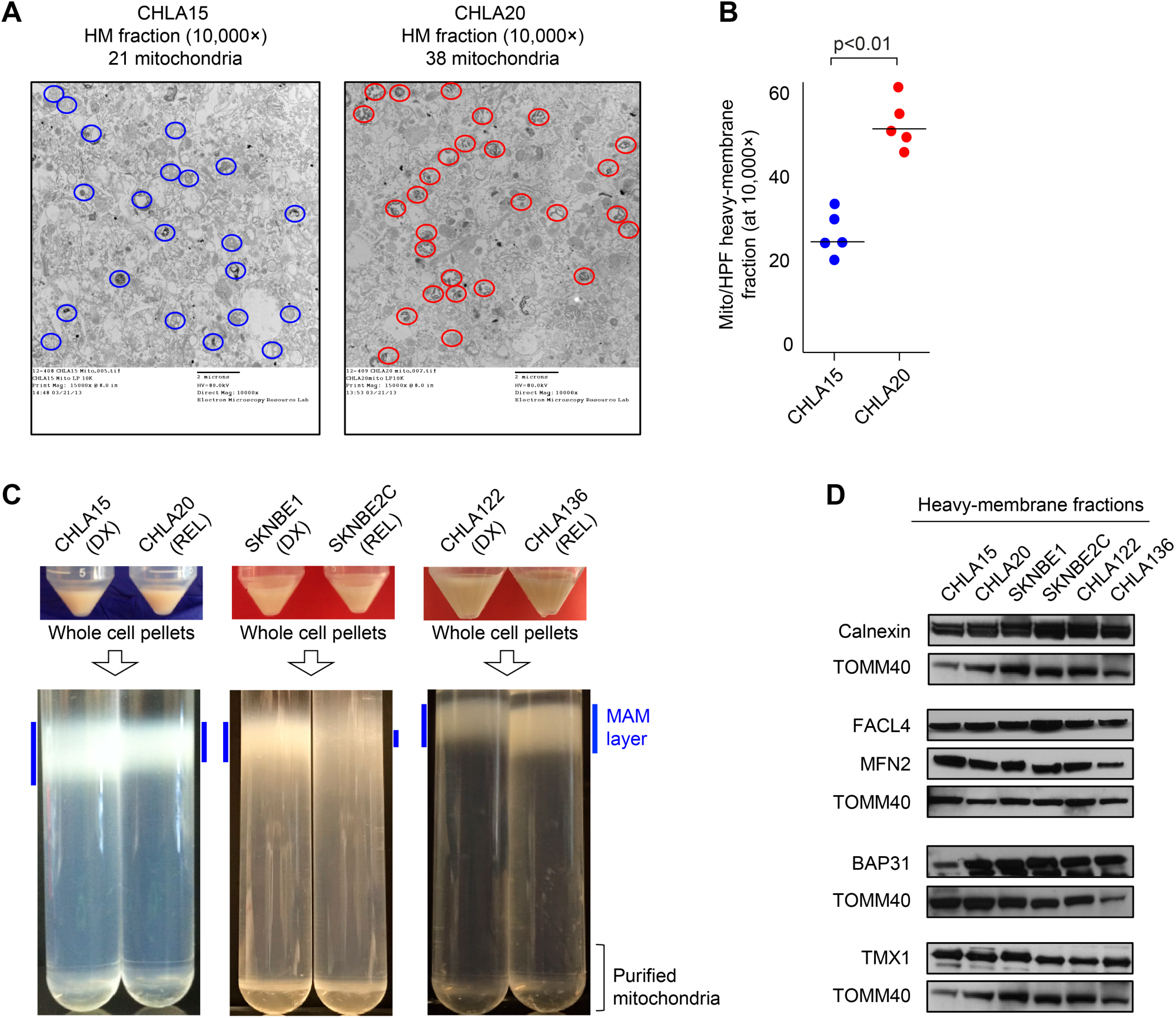
Reduced MAM content in neuroblastomas obtained at the time of REL in comparison to DX. **(A)** Electron micrographs images of heavy-membrane mitochondria-enriched fractions from CHLA15 and CHLA20 cells identified more mitochondria per volume in REL CHLA20 cells, graphically represented in **(B**); statistical analyses performed using an unpaired two-sided Student’s t-test. **(C)** Heavy-membrane fractions obtained from equal cell input were separated by 30% Percoll gradient ultracentrifugation into MAM fractions (highlighted by blue-bars) and purified mitochondria for three DX/REL pairs. **(D)** Immunoblot detection of various MAM-and ER-localized proteins from heavy-membrane mitochondria-enriched lysates in DX and REL tumors, with TOMM40 as a loading control for mitochondrial protein input.

**Supplemental Figure 6.**
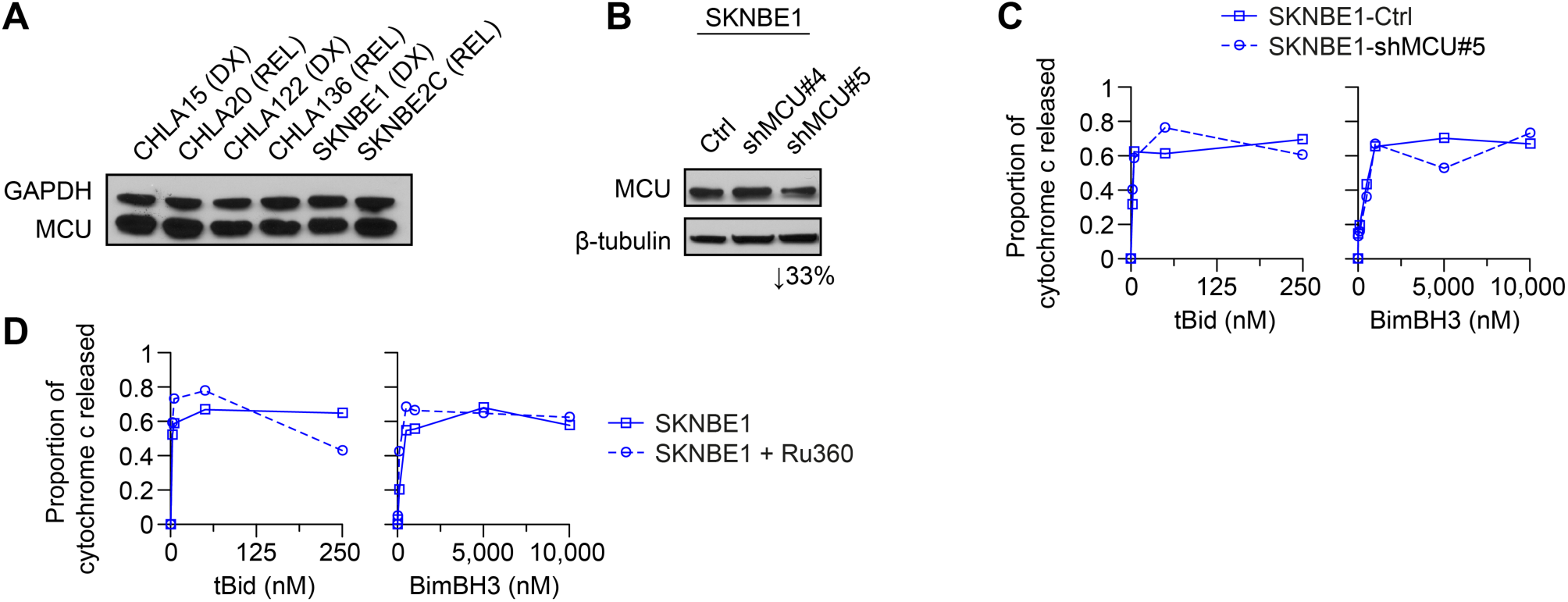
Attenuating transfer of calcium from ER to mitochondria does not attenuate mitochondrial stress responses. **(A)** Immunoblot detection of MCU (mitochondrial calcium uniporter) expression across patient-matched pairs of at-diagnosis (DX) and post-relapse (REL) neuroblastomas. **(B)** Silencing of the mitochondrial calcium uniporter (MCU) in SKNBE1 cells, with ∼33% reduction in MCU at the protein level in SKNBE1-shMCU#5. **(C)** Mitochondrial cytochrome c release for SKNBE1-shMCU#5 cells following exposure to tBid or BimBH3 peptide. **(D)** Mitochondrial cytochrome c release for SKNBE1 cells following exposure to tBid or BimBH3 peptide, with or without 30-minute Ruthenium-360 pre-treatment (Ru360). For C and D, data points are mean of duplicate wells (SD<0.05 at all points) in a representative experiment from at least three biological replicates.

## REFERENCES

1. Holohan C, Van Schaeybroeck S, Longley DB, and Johnston PG. Cancer drug resistance: an evolving paradigm. Nature reviews Cancer. 2013;13(10):714–26.

2. Olivier M, Hollstein M, and Hainaut P. TP53 mutations in human cancers: origins, consequences, and clinical use. Cold Spring Harb Perspect Biol. 2010;2(1):a001008.

3. Hugo W, et al. Non-genomic and Immune Evolution of Melanoma Acquiring MAPKi Resistance. Cell. 2015;162(6):1271–85.

4. Katayama R, et al. Mechanisms of acquired crizotinib resistance in ALK-rearranged lung Cancers. Sci Transl Med. 2012;4(120):120ra17.

5. Wilson FH, et al. A functional landscape of resistance to ALK inhibition in lung cancer. Cancer Cell. 2015;27(3):397–408.

6. Chong CR, and Janne PA. The quest to overcome resistance to EGFR-targeted therapies in cancer. Nat Med. 2013;19(11):1389–400.

7. Lord CJ, and Ashworth A. Mechanisms of resistance to therapies targeting BRCA-mutant cancers. Nat Med. 2013;19(11):1381–8.

8. Lito P, Rosen N, and Solit DB. Tumor adaptation and resistance to RAF inhibitors. Nat Med. 2013;19(11):1401–9.

9. Pinto NR, et al. Advances in Risk Classification and Treatment Strategies for Neuroblastoma. J Clin Oncol. 2015;33(27):3008–17.

10. Keshelava N, Seeger RC, Groshen S, and Reynolds CP. Drug resistance patterns of human neuroblastoma cell lines derived from patients at different phases of therapy. Cancer Res. 1998;58(23):5396–405.

11. Keshelava N, Seeger RC, and Reynolds CP. Drug resistance in human neuroblastoma cell lines correlates with clinical therapy. Eur J Cancer. 1997;33(12):2002–6.

12. Matthay KK, et al. Long-term results for children with high-risk neuroblastoma treated on a randomized trial of myeloablative therapy followed by 13-cis-retinoic acid: a children’s oncology group study. J Clin Oncol. 2009;27(7):1007–13.

13. Kim H, et al. Hierarchical regulation of mitochondrion-dependent apoptosis by BCL-2 subfamilies. Nat Cell Biol. 2006;8(12):1348–58.

14. Ren D, et al. BID, BIM, and PUMA are essential for activation of the BAX-and BAK-dependent cell death program. Science. 2010;330(6009):1390–3.

15. Sarosiek KA, et al. BID preferentially activates BAK while BIM preferentially activates BAX, affecting chemotherapy response. Mol Cell. 2013;51(6):751–65.

16. Tan TT, et al. Key roles of BIM-driven apoptosis in epithelial tumors and rational chemotherapy. Cancer Cell. 2005;7(3):227–38.

17. Csordas G, Weaver D, and Hajnoczky G. Endoplasmic Reticulum-Mitochondrial Contactology: Structure and Signaling Functions. Trends Cell Biol. 2018;28(7):523–40.

18. Rowland AA, and Voeltz GK. Endoplasmic reticulum-mitochondria contacts: function of the junction. Nat Rev Mol Cell Biol. 2012;13(10):607–25.

19. Arruda AP, Pers BM, Parlakgul G, Guney E, Inouye K, and Hotamisligil GS. Chronic enrichment of hepatic endoplasmic reticulum-mitochondria contact leads to mitochondrial dysfunction in obesity. Nat Med. 2014;20(12):1427–35.

20. Cali T, Ottolini D, and Brini M. Calcium and endoplasmic reticulum-mitochondria tethering in neurodegeneration. DNA Cell Biol. 2013;32(4):140–6.

21. Hedskog L, et al. Modulation of the endoplasmic reticulum-mitochondria interface in Alzheimer’s disease and related models. Proc Natl Acad Sci U S A. 2013;110(19):7916–21.

22. Area-Gomez E, et al. Upregulated function of mitochondria-associated ER membranes in Alzheimer disease. The EMBO journal. 2012;31(21):4106–23.

23. Deng J, Carlson N, Takeyama K, Dal Cin P, Shipp M, and Letai A. BH3 profiling identifies three distinct classes of apoptotic blocks to predict response to ABT-737 and conventional chemotherapeutic agents. Cancer Cell. 2007;12(2):171–85.

24. Gavathiotis E, et al. BAX activation is initiated at a novel interaction site. Nature. 2008;455(7216):1076–81.

25. Goldsmith KC, et al. Mitochondrial Bcl-2 family dynamics define therapy response and resistance in neuroblastoma. Cancer Res. 2012;72(10):2565–77.

26. Sarosiek KA, Ni Chonghaile T, and Letai A. Mitochondria: gatekeepers of response to chemotherapy. Trends Cell Biol. 2013;23(12):612–9.

27. Bresler SC, et al. ALK mutations confer differential oncogenic activation and sensitivity to ALK inhibition therapy in neuroblastoma. Cancer Cell. 2014;26(5):682–94.

28. Park JH, Zhuang J, Li J, and Hwang PM. p53 as guardian of the mitochondrial genome. FEBS Lett. 2016;590(7):924–34.

29. Annunziata I, Patterson A, and d’Azzo A. Mitochondria-associated ER membranes (MAMs) and glycosphingolipid enriched microdomains (GEMs): isolation from mouse brain. J Vis Exp. 2013(73):e50215.

30. Chipuk JE, et al. Sphingolipid metabolism cooperates with BAK and BAX to promote the mitochondrial pathway of apoptosis. Cell. 2012;148(5):988–1000.

31. Hajnoczky G, Csordas G, Madesh M, and Pacher P. Control of apoptosis by IP(3) and ryanodine receptor driven calcium signals. Cell Calcium. 2000;28(5-6):349–63.

32. Rizzuto R, and Pozzan T. Microdomains of intracellular Ca2+: molecular determinants and functional consequences. Physiol Rev. 2006;86(1):369–408.

33. Rimessi A, Giorgi C, Pinton P, and Rizzuto R. The versatility of mitochondrial calcium signals: from stimulation of cell metabolism to induction of cell death. Biochim Biophys Acta. 2008;1777(7-8):808–16.

34. Csordas G, et al. Structural and functional features and significance of the physical linkage between ER and mitochondria. The Journal of cell biology. 2006;174(7):915–21.

35. Baughman JM, et al. Integrative genomics identifies MCU as an essential component of the mitochondrial calcium uniporter. Nature. 2011;476(7360):341–5.

36. De Stefani D, Raffaello A, Teardo E, Szabo I, and Rizzuto R. A forty-kilodalton protein of the inner membrane is the mitochondrial calcium uniporter. Nature. 2011;476(7360):336–40.

37. de Brito OM, and Scorrano L. Mitofusin 2 tethers endoplasmic reticulum to mitochondria. Nature. 2008;456(7222):605–10.

38. Moulis M, Grousset E, Faccini J, Richetin K, Thomas G, and Vindis C. The Multifunctional Sorting Protein PACS-2 Controls Mitophagosome Formation in Human Vascular Smooth Muscle Cells through Mitochondria-ER Contact Sites. Cells. 2019;8(6).

39. Simmen T, et al. PACS-2 controls endoplasmic reticulum-mitochondria communication and Bid-mediated apoptosis. The EMBO journal. 2005;24(4):717–29.

40. Konieczkowski DJ, Johannessen CM, and Garraway LA. A Convergence-Based Framework for Cancer Drug Resistance. Cancer Cell. 2018;33(5):801–15.

41. Danial NN, and Korsmeyer SJ. Cell death: critical control points. Cell. 2004;116(2):205–19.

42. Montero J, et al. Drug-induced death signaling strategy rapidly predicts cancer response to chemotherapy. Cell. 2015;160(5):977–89.

43. Ni Chonghaile T, et al. Pretreatment mitochondrial priming correlates with clinical response to cytotoxic chemotherapy. Science. 2011;334(6059):1129–33.

44. Vo TT, et al. Relative mitochondrial priming of myeloblasts and normal HSCs determines chemotherapeutic success in AML. Cell. 2012;151(2):344–55.

45. Hata AN, Engelman JA, and Faber AC. The BCL2 Family: Key Mediators of the Apoptotic Response to Targeted Anticancer Therapeutics. Cancer Discov. 2015;5(5):475–87.

46. Uccini S, et al. Morphological and molecular assessment of apoptotic mechanisms in peripheral neuroblastic tumours. Br J Cancer. 2006;95(1):49–55.

47. Hinds MG, et al. Bim, Bad and Bmf: intrinsically unstructured BH3-only proteins that undergo a localized conformational change upon binding to prosurvival Bcl-2 targets. Cell Death Differ. 2007;14(1):128–36.

48. Walensky LD, et al. A stapled BID BH3 helix directly binds and activates BAX. Mol Cell. 2006;24(2):199–210.

49. Kale J, Osterlund EJ, and Andrews DW. BCL-2 family proteins: changing partners in the dance towards death. Cell Death Differ. 2018;25(1):65–80.

50. Hoppins S, and Nunnari J. Cell Biology. Mitochondrial dynamics and apoptosis--the ER connection. Science. 2012;337(6098):1052–4.

51. Renault TT, et al. Mitochondrial shape governs BAX-induced membrane permeabilization and apoptosis. Mol Cell. 2015;57(1):69–82.

52. Tait SW, et al. Resistance to caspase-independent cell death requires persistence of intact mitochondria. Dev Cell. 2010;18(5):802–13.

53. Howatson AF, and Ham AW. Electron microscope study of sections of two rat liver tumors. Cancer Res. 1955;15(1):62–9.

54. Scorrano L, et al. BAX and BAK regulation of endoplasmic reticulum Ca2+: a control point for apoptosis. Science. 2003;300(5616):135–9.

55. Patergnani S, et al. The endoplasmic reticulum mitochondrial calcium cross talk is downregulated in malignant pleural mesothelioma cells and plays a critical role in apoptosis inhibition. Oncotarget. 2015;6(27):23427–44.

56. Csordas G, et al. Imaging interorganelle contacts and local calcium dynamics at the ER-mitochondrial interface. Mol Cell. 2010;39(1):121–32.

57. Martino Adami PV, et al. Perturbed mitochondria-ER contacts in live neurons that model the amyloid pathology of Alzheimer’s disease. J Cell Sci. 2019;132(20).

58. Schauder CM, et al. Structure of a lipid-bound extended synaptotagmin indicates a role in lipid transfer. Nature. 2014;510(7506):552–5.

59. Renault TT, and Chipuk JE. Death upon a kiss: mitochondrial outer membrane composition and organelle communication govern sensitivity to BAK/BAX-dependent apoptosis. Chemistry & biology. 2014;21(1):114–23.

60. Paillard M, et al. Depressing mitochondria-reticulum interactions protects cardiomyocytes from lethal hypoxia-reoxygenation injury. Circulation. 2013;128(14):1555–65.

61. Esterberg R, Hailey DW, Rubel EW, and Raible DW. ER-mitochondrial calcium flow underlies vulnerability of mechanosensory hair cells to damage. The Journal of neuroscience : the official journal of the Society for Neuroscience. 2014;34(29):9703–19.

62. Ottolini D, Cali T, Negro A, and Brini M. The Parkinson disease-related protein DJ-1 counteracts mitochondrial impairment induced by the tumour suppressor protein p53 by enhancing endoplasmic reticulum-mitochondria tethering. Hum Mol Genet. 2013;22(11):2152–68.

63. Zampese E, Fasolato C, Kipanyula MJ, Bortolozzi M, Pozzan T, and Pizzo P. Presenilin 2 modulates endoplasmic reticulum (ER)-mitochondria interactions and Ca2+ cross-talk. Proc Natl Acad Sci U S A. 2011;108(7):2777–82.

64. Rieusset J. Mitochondria and endoplasmic reticulum: mitochondria-endoplasmic reticulum interplay in type 2 diabetes pathophysiology. The international journal of biochemistry & cell biology. 2011;43(9):1257–62.

65. Bartok A, et al. IP3 receptor isoforms differently regulate ER-mitochondrial contacts and local calcium transfer. Nat Commun. 2019;10(1):3726.

66. Tosatto A, et al. The mitochondrial calcium uniporter regulates breast cancer progression via HIF-1alpha. EMBO Mol Med. 2016;8(5):569–85.

67. Csordas G, and Hajnoczky G. Sorting of calcium signals at the junctions of endoplasmic reticulum and mitochondria. Cell Calcium. 2001;29(4):249–62.

